# Intestinal epithelial differentiation factor HNF4α restrains homeostatic interferon signaling and is required for its full induction during *Salmonella* infection

**DOI:** 10.64898/2026.11.14.766693

**Authors:** Courtney Rivet-Noor, Colin R. Lickwar, Matt C. Tillman, Lauren M. Orr, Whitney L. M. DeLoach, Keir M. Balla, Faith V. You, Rebecca L. Bacon, Emma Heckenburg, Carlton W. Anderson, Sarah E. Blutt, Carolyn B. Coyne, Neeraj K. Surana, Stacy M. Horner, John F. Rawls

## Abstract

Intestinal epithelial cells (IECs) must interpret microbial signals to transcriptionally adapt their identity and function. Which transcription factors (TFs) mediate those adaptations, and whether competing transcriptional pathways signal within the same or in different IEC subpopulations, remains unclear. We find that microbiota colonization shifts nearly all IECs toward an interferon-driven inflammatory program at the expense of genes involved in enterocyte differentiation and metabolism. These opposing gene expression programs are associated with different TF motifs and are inversely correlated within individual cells. We discover that HNF4α, a TF central to enterocyte differentiation, acts as a brake on interferon signaling *in vitro*. In accord, loss of HNF4α in IECs in mice elevates baseline interferon-stimulated gene expression but prevents its full induction during infection with the bacterial pathogen *Salmonella*. Together our findings reveal a novel mechanism by which opposing intestinal epithelial differentiation and interferon signaling programs interact within the same cell to maintain proper transcriptional control and homeostasis.

## Introduction

Animal cells use their shared genome to differentiate into specialized types, yet those cells must also sense and respond to the microbial world. Cell identity and microbial responsiveness are often governed by distinct signaling pathways and downstream transcription factors (TFs), but how these programs are coordinated within individual cells to balance tissue identity against microbial response remains poorly understood (1, 2).

This balance is especially consequential in the intestinal epithelium which is comprised of absorptive enterocytes and multiple secretory lineages arising from intestinal stem cells (3). Enterocytes absorb dietary nutrients and form the physical barrier separating the body from the bacteria, fungi, and viruses residing in the lumen (4). As enterocytes and other intestinal epithelial cells (IECs) migrate up the crypt-villus axis, developmental signals such as Bone morphogenetic proteins (BMPs) generate spatially organized (“zonated”) cell states that differ in absorptive capacity (5–9). IECs are also continually exposed to microbial products and to cytokines induced by the microbiota (10–13). For example, microbe associated molecular patterns induce production of interferons (IFNs) by both immune cells and IECs themselves, and these IFNs signal in autocrine and paracrine fashions through JAK/STAT pathways to induce IFN-stimulated genes (ISGs) that support antimicrobial defense (10–12, 14–18). How the pathways that mediate microbial and IFN responses interact with those that maintain differentiated intestinal epithelial physiology remains unknown.

Hepatocyte nuclear factor 4 alpha (HNF4α; NR2A1) is a nuclear receptor TF that promotes epithelial differentiation in the intestine and kidney (19, 20). In the intestine it drives brush border formation, absorptive enterocyte differentiation, and fatty acid oxidation by acting with its paralog HNF4γ and BMP signaling (20–22). These roles for HNF4α in intestinal identity appear to be well conserved across bilaterian animals (23–26). HNF4α is also relevant to intestinal health: IEC-specific deletion in mice causes spontaneous colitis (27, 28) and increases susceptibility to DSS-induced colitis (29–31). The human inflammatory bowel diseases ulcerative colitis and Crohn’s disease have also both been associated with *HNF4A* variants (32–34) and significant reductions in *HNF4A* levels (31).

We recently discovered that HNF4α is also involved in the host response to microbes. Microbiota colonization broadly suppresses HNF4α chromatin occupancy and HNF4α-driven transcription in jejunal IECs (24, 25, 35), while coincidently inducing inflammatory programs (12, 24, 36) enriched for IRF, STAT, and ETS motifs (24, 37). These programs are inversely correlated (24, 37, 38), and deleting HNF4α in mouse IECs increases inflammatory gene expression (27, 28). Together, these findings suggest that HNF4α-driven and inflammatory programs oppose one another. However, this much of this evidence comes from bulk IEC studies (24, 35, 37), which cannot distinguish whether individual cells shift between the two competing programs or if the tissue changes in its proportion of cell types separately expressing those programs. Further, whether the two programs interact within single cells, and whether that interaction affects the response to pathogen infection and microbiota colonization, is unknown.

Here, we asked whether HNF4α functionally interacts with IFN signaling. We profiled jejunal IECs from germ-free (GF) and conventionalized (CV) mice by single-cell RNA sequencing (scRNA-seq). We found that conventionalization induced an IFN-related program across all IEC subpopulations, with HNF4α motifs enriched among microbially down-regulated genes and IRF, STAT, and NF-κB motifs enriched among up-regulated genes. Within single cells, HNF4α-associated and inflammatory gene expression programs were inversely correlated. Moving to HEK293T cells to facilitate pathway manipulations, we found HNF4α suppressed microbial stimulation of IFN and NF-κB activity, an effect that required its DNA-binding domain and the induction of multiple secondary mediators. Validation studies confirmed that loss of HNF4α in IECs elevated baseline ISG expression in organoids and *in vivo*. During pathogenic *Salmonella* infection, however, HNF4α-deficient mice failed to fully induce ISGs and other terminally differentiated enterocyte genes, suggesting HNF4α function may be required for maximal ISG induction. These findings are important because they extend HNF4α’s role beyond enterocyte differentiation to restraining epithelial ISG induction and suggest that the balance between metabolic identity and IFN response shapes how the epithelium responds to infection.

## Results

### Response to microbes in jejunal enterocyte by scRNA-seq

To identify transcriptional and cell type changes induced by microbiota colonization, we isolated EpCAM+ intestinal epithelial cells (IECs) from the jejunum of adult mice reared germ-free (GF) and ex-GF mice colonized for 2 weeks with a fecal microbiota from specific pathogen-free mice (conventionalized, CV) (Figure 1A,B and Supplemental Figure 1A-G). Processing through Cell Ranger and Seurat pipelines identified 9114K IECs (4481K CV and 4633K GF) that were reduced to 6442K IECs (3244 CV and 3198 GF) following filtering by feature and mitochondrial thresholds (Figures 1B and Supplemental Figure1A). As expected, the dataset was predominated by enterocytes (Clusters 0-6) stratified by small intestinal region along UMAP 1 (Figures 1C, Supplemental Figure 2A + Supplemental Table 1) (4, 43). This revealed a subset of ileocyte type cells (*Fabp6* and *Slc10a2*, Cluster 4; Figure 1C). While small intestinal (SI) regions are not typically based on distinct anatomical landmarks, recent high-resolution regional intestinal analysis has shown, dispersed within any of the major SI regions, gradients of cells expressing regional markers typically associated with neighboring regions (43). Along UMAP 2, cell clusters corresponded with progressive bands of the major differentiation markers for the enterocyte villus base (*Reg1*, Cluster 0), villus middle (*Slc2a2*, Clusters 0,1,2,3,4,6) and villus tip (*Ada*, Cluster 5). A small portion of secretory cells including goblet cells, EECs (*Agr2*, *Neurod1*; Cluster 8), and Paneth cells (*Lyz1*; Cluster 9) were also identified (Figure 1C and Supplemental Figure 2B). While of interest, these cell populations were not large enough to facilitate extensive additional subsetting or analysis, but could be interesting targets for future study.

**Figure 1:**
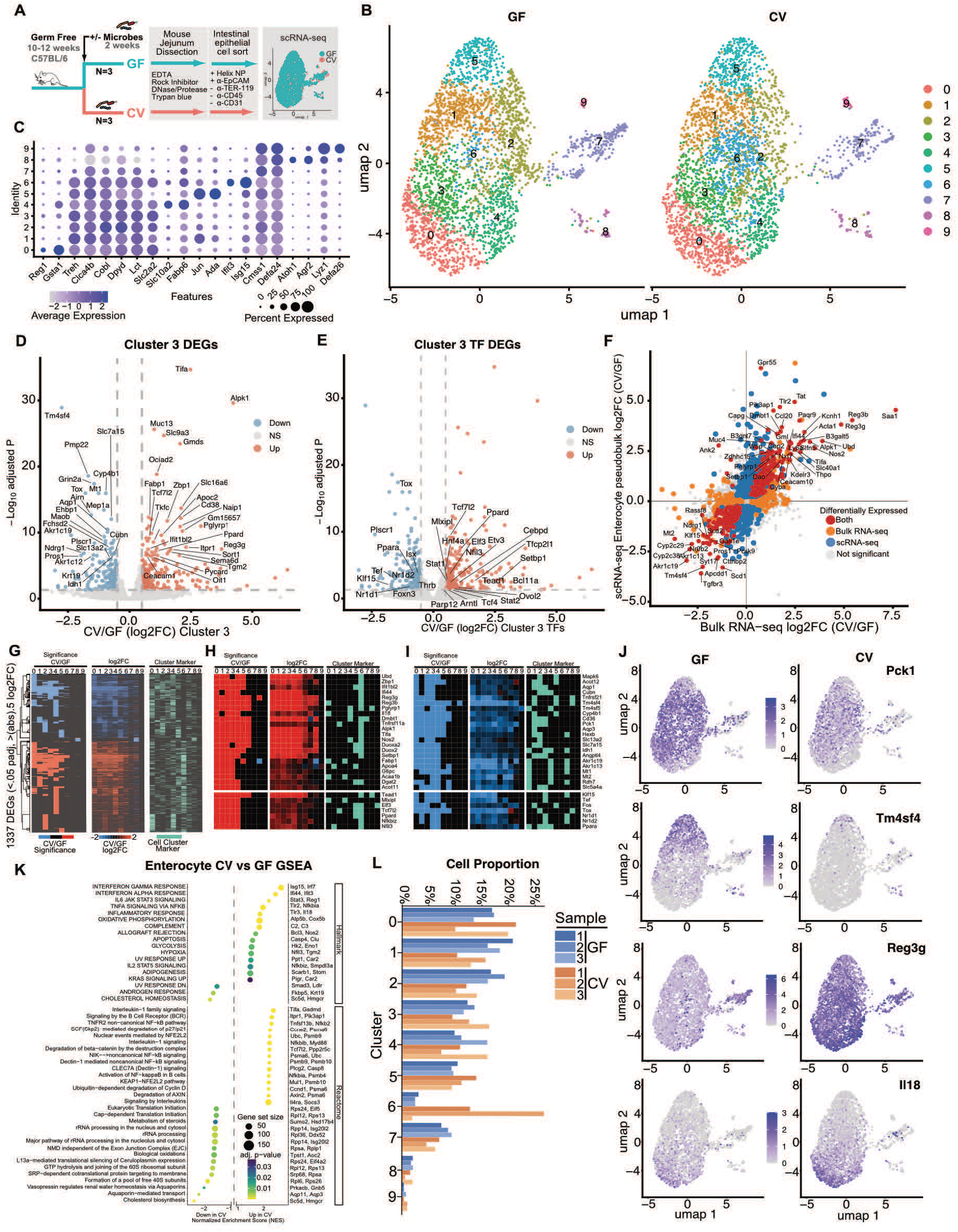
scRNA-seq of cell type specific microbial responses in jejunal mouse IECs. **(A)** Experimental schematic of scRNA-seq in CV and GF mouse jejunal IECs. **(B)** UMAP of all jejunal IECs in GF and CV. **(C)** Dotplot of cell cluster markers across enterocytes, secretory, and proliferative cells. **(D)** Volcano plot of significantly differential expressed genes (CV/GF) using cluster 3 as a representative for all genes and **(E)** restricted to transcription factors (TFs). **(F)** Scatterplot comparing previously published gene expression changes in bulk jejunal IECs (CV/GF) (35)to a computational combination of all enterocytes from this study (Clusters 0-6, CV/GF). **(G)** Genes significantly different in at least one cluster comparison (CV/GF) hierarchically clustered by call and colored for up, down, or no significance for each cluster for (CV/GF, left), continuous log_2_FC (CV/GF) values for each scRNA-seq cluster (middle), and if the gene is called as a cluster marker (right). Example genes that are **(H)** upregulated or **(I)** downregulated in CV/GF subset from heatmaps in (G). **(J)** FeaturePlots of cell level expression values of example genes in the GF (left) and CV condition (right) that have expression generally restricted to a subset of enterocytes. **(K)** GSEA analysis of CV/GF comparison for grouped scRNA-seq enterocyte clusters displaying a subset of the most significant terms. Example genes for each significant term are listed on the right. **(L)** Barplot of cell proportion per biological replicate sample separated by cluster and condition.

### Microbial responses are generally distributed across enterocyte subtypes

Pseudobulk differential expression analysis of each cluster’s microbial responses (CV/GF) identified numerous canonical responsive upregulated (*Reg3g*, *Tifa*) and downregulated genes (*Tm4sf4*, *Akr1c19*, *Mt1*), including at representative enterocyte cluster 3 (Figure 1D + Supplemental Table 2). Similarly, transcription factors (TFs) like *Elf3*, *Nfil3*, *Stat1/2* were upregulated and nuclear receptor (*Ppara*, *Nr1d1/2*, *Thrb*) and bZip TFs (*Dbp*, *Tef*) were downregulated (Figure 1E). Accordingly, log_2_FC for grouped enterocyte clusters’ microbial response was highly concordant with the microbial response of bulk jejunum enriched for IECs (Figure 1F) (24, 35). We were curious to identify cluster-specific microbial responses, so we hierarchically clustered all genes that were differential in any scRNA-seq cluster in CV/GF (Figure 1G). While we identified upregulated (Figure 1H) and downregulated (Figure 1I) significant genes that were restricted to certain scRNA-seq clusters, we were interested to see that the consistent directionality of the fold change was almost always broadly shared in additional clusters, even if not significant in those clusters (Figure 1G-I + Supplemental Figure 2C-F). Additionally, essentially no genes were coincidently and significantly differentially expressed in the opposite direction in any clusters. As a result, though many genes’ expression patterns were highly specific to an enterocyte cluster, it was difficult to find examples where the microbial response was similarly cluster restricted (Figures 1G-J). This implies that expression gradients of enterocyte differentiation and regionality interact with microbial responsive programs diffusely, and that microbially responsive programs are not highly cluster restrictive nor insulated transcriptionally.

Gene set enrichment analysis (GSEA) of grouped enterocyte cell clusters identified strong enrichment of inflammation-related terms including interferon response (*Isg15*, *Irf6*, *Ifi44*), IL6/Jak/Stat (*Stat3*, *Reg1*), interleukin (*Tifa*), and NF-κB (*Nfkb2, Nfkbia*) signaling in genes that were upregulated upon colonization. Downregulated terms included those associated with cholesterol (*Sc5d*, *Hmgcr*) and steroid metabolism (*Sumo2*, *Hsd17b4*), translation (*Rps24*, *Eif5*), rRNA processing (*Rpp14*, *Isg20l2*), and Aquaporin-mediated transport (*Aqp11*, *Aqp3*) (Figure 1K).

We considered that shifts in complex transcriptional programs or cell differentiation pathways in response to microbiota may reflect transcriptional changes amongst like-cells or differences in enterocyte subtype proportion. We therefore looked for significant differences in cell cluster proportion across the CV and GF conditions. While no clusters were significantly different across conditions using this experimental design and sequencing depth, we found a 1.4-fold decrease in cell proportion in Cluster 2 between CV and GF and a >5-fold increase in Cluster 6, though this was driven by two of three replicates (Figure 1L and Supplemental Figure 2G-H).

### Interferon stimulated genes are highly expressed as Cluster 6 markers and correlate with microbial responsive genes

Recently, foci of epithelial cells expressing interferon simulated genes (ISG) have been found on small intestinal villi, though their regulation and genesis are not fully understood (12, 39). We found that enterocyte Cluster 6 is enriched for high ISG and inflammatory gene expression, which was also induced transcriptionally in the surrounding clusters (Figures 1H and 2A,B). Although transcripts for interferon genes themselves were not a part of this Cluster 6 signature, this may be attributable to the known transience, sparsity, and stochasticity of IFN gene induction (40–43). While there is apparent variability in Cluster 6 cell number induction (Figure 1L), importantly, ISG-high cells exist in the GF condition suggesting that microbes are not absolutely necessary for these cells.

**Figure 2:**
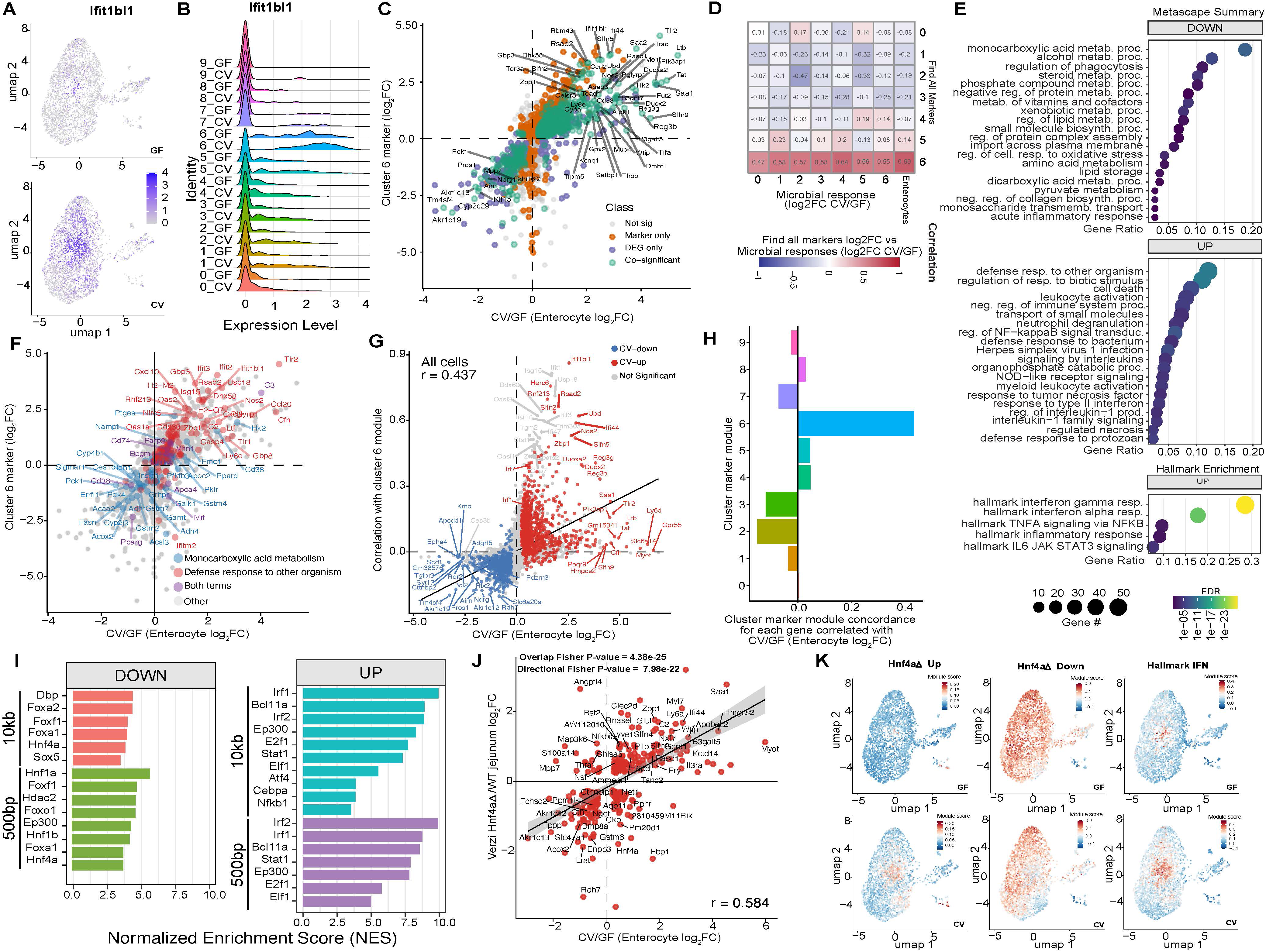
Following colonization by microbes, enterocytes reduce their identity and adopt an ISG program that is concentrated in a subset of IECs and may be suppressed by HNF4α. **(A)** Featureplot of ISG *Ifit1bl1* in GF and CV conditions. **(B)** Ridgeplot of ISG *Ifit1bl1* highlighting its broad microbial induction and concentrated expression in cluster 6. **(C)** Scatterplot comparing scRNA-seq microbial response log_2_FC (CV/GF) to each gene’s Cluster 6 find all markers log_2_FC value colored by significance categories. High log_2_FC represents genes specifically expressed in cluster 6 relative to all other clusters. **(D)** Heatmap of linear correlation between cluster-specific microbial response log_2_FC and enterocyte cluster-specific find all marker log_2_FC as in 2C. Only markers for cluster 6 are also broadly induced by microbes. **(E)** Metascape analysis for Down and Up gene groups significant as cluster markers and microbially responsive in cluster 6. **(F)** Scatterplot as in (C) colored by genes that are members of enriched GO terms for Down and Up gene lists. **(G)** Scatterplot of cell-by-cell correlation of a cluster 6 marker gene module score against all other genes and colored by microbial responsiveness in enterocyte clusters as a group. **(H)** Correlation as defined in (G) for all clusters marker gene modules cell by cell expression correlation for all genes relative to microbial response (Enterocyte log_2_FC (CV/GF)). **(I)** Motif enrichment for a 10kb (top) or 500bp (bottom) region surrounding genes in the Down (left) or Up (right) gene groups. **(J)** Scatterplot of microbial response log_2_FC (CV/GF) versus effect of HNF4α loss log_2_FC (HNF4α KO/WT)(47). **(K)** Featureplots of HNF4α KO Up significant genes (left), HNF4α KO Down significant genes (middle), and Hallmark interferon response genes (right) split by the GF (top) or CV (bottom) condition.

We considered the possibility that viruses might contribute to ISG induction in the GF state or further induce ISGs in CV. PCR assays for a panel of mouse viruses using fecal pellets from the colonies of GF and specific pathogen-free (SPF) mice used in this study were all negative (not shown). We pursued this further by screening our scRNA-seq data for viral transcripts to test the possibility that the ISG responses observed in CV or GF mice were driven by cryptic viral infections. We de novo assembled approximately 60K transcripts from CV and GF samples and identified 119 transcripts with high sequence similarity to endogenous retroviruses (ERVs) in the mouse genome (Supplemental Figure 3A,B + Supplemental Tables 3, 4). We did not identify transcripts with sequence similarity to exogenous viruses that infect vertebrates and thus did not identify evidence for cryptic infections in CV or GF mice. Quantification of ERV and transposable element (TE) transcripts revealed differential expression of individual loci between CV and GF mice but no net difference in the total ERV/TE load (Supplemental Figure 3B + Supplemental Tables 3, 4). These results argue against a link between ERV/TE expression and the ISG signature we observed. This leads us to suggest direct, tonic IFN or microbial sensing as the best explanation for the ISG-high states in GF and CV mice, respectively.

To interrogate if induction of the Cluster 6 program was coincident with microbially regulated genes, we compared Cluster 6 marker log_2_FC enrichment with microbial response across enterocytes. We found impressive linear correlation with both microbially-induced and microbially-suppressed genes and Cluster 6 marker gene scoring (Figure 2C). Importantly, this strong positive correlation was only found when using marker enrichment from Cluster 6 (Figure 2D). We identified two salient groups of genes: those that were positive Cluster 6 marker genes and induced by microbes (Up), and those that were negative Cluster 6 marker genes and suppressed by microbes (Down). GO term analysis identified broad IFN and response to bacterium responses in the Up group and metabolic programs in the Down group, suggesting these programs may be reciprocally regulated in the intestine (44) (Figure 2E,F). Comparing each gene’s expression relative to other genes or programs within each cell should identify whether these Up and Down programs are, in fact, juxtaposed within cells or if their programs are just discordant on average. We therefore looked at the linear correlation between the top 50 Cluster 6 marker genes’ expression and every other gene’s expression across all cells and found a strong linear correlation (Figure 2G). This implies ISG-high expressing cells are more likely to have strong repression of microbially-suppressed genes and vice-versa. This property is specific to Cluster 6 markers, is not due solely to sparse or non-uniform sequencing (Figure 2H and Supplemental Figure 4A-C), and this relationship increases as more cells are considered (Supplemental Figure 4D-L). Notably, while the Up group was dominated by ISG genes, it also included non-ISG inflammatory genes like *Reg3g, Reg3b, Duox2, Duoxa2, Saa1,* and *Tlr2* (Figure 2F,G). Together, this suggests that following exposure to microbes, most GF enterocytes are replenished by those with reduced general tissue identity in favor of amplifying an ISG and inflammation dominant program (Figure 2D).

### HNF4 is associated with microbially-down and IRF, Stat, NF-κB, and Ets microbially-up genes

We next looked for enrichment of transcription factor binding site (TFBS) motifs in our Up and Down groups that showed correlation between microbial response and Cluster 6 identity (Figure 2C). We found that DBP, HNF4, and Fox factor TF motif enrichments are associated with the Down gene group and IRF, Stat, NF-κB are potential large-scale regulators of the Up program (Figure 2I). We and others have previously implicated these pathways in the microbial response (44). However, our scRNA-seq implies these programs may directly antagonize each other within individual cells, suggesting this antagonism is a critical intersection balancing metabolic and inflammatory function.

### HNF4 -regulated genes correlate with reciprocally expressed microbially-regulated genes

We have previously shown that a majority of microbially suppressed genes are positively regulated by Hnf4 in the zebrafish digestive tract (24). Similar analysis in mouse showed that IEC-specific HNF4 deletion results in reduction of microbially suppressed genes and induction of microbially induced genes, though only suppressed genes appear to be direct HNF4 targets (35). While HNF4 and HNF4γ show substantial redundancy and deletion of both genes results in profound loss of enterocyte identity (22, 25), IEC-specific HNF4 deletion frequently results in reduction of enterocyte differentiation genes and can lead to sporadic intestinal inflammation (27, 45, 46). Importantly, GF mice harboring the same IEC-specific HNF4 deletion develop little or no inflammation, suggesting loss of HNF4 may not always derepress inflammation signaling, but may predispose cells to inflammatory signaling depending on environment or context (27). Our microbially regulated scRNA-seq genes also correlate with genes regulated by HNF4 (Figure 2J). This included microbially suppressed genes being HNF4 activated (reduced on HNF4 loss; *Rdh7*, *Acox2*, *Akr1c13*), and microbially activated genes being HNF4 suppressed (increased on HNF4 loss; *Ifi44*, *Zbp1*, *Nfkbia*, *Ly6a*) and, as a result, coincident with expression of ISGs in Cluster 6 (Figure 2J-K) (47). IEC activation of IFN signaling and Cluster 6 markers occurs in multiple independent IEC-specific HNF4 deletion datasets (Supplemental Figure 4M-N), though this response may not be fully penetrant, consistent with sporadic impact of HNF4 loss (35). The mechanism for the apparent HNF4 - influenced interaction between microbially induced ISGs and microbially suppressed metabolic genes is not known.

### HNF4 suppresses interferon and NF-κB pathways *in vitro*

Our single-cell RNA-seq and HNF4 deletion dataset analysis suggested that HNF4 - driven and ISG-high programs (enriched for IRF, STAT, and NF-κB TFBS motifs) oppose one another and are likely to interact within the same cell (Figure 2G,I). However, direct manipulation of signaling cascades and TF pathways *in vivo* poses significant technical challenges. To better address the hypothesized interactions between HNF4 and ISG programs, we moved to a more reductionist system in HEK293T cells.

Both *in vivo* and *in vitro*, microbes and their products can stimulate the cellular “pathogen-sensing” arm and initiate signaling cascades (RIG-I, TLR4, etc) to promote NF-κB and IFN production (Figure 3A) (10, 11, 48–53). Once produced, NF-κB and IFN can act within the cell to influence the production of ISGs. NF-κB binds IFN enhanceosomes to recruit gene inducing co-factors and further promoter IFN signaling, while IFNs signal in an autocrine or paracrine manner to activate the “IFN amplification arm” via the JAK/STAT pathway (54, 55). The IFN amplification arm in turn, induces ISG expression through interferon-stimulated response elements (ISRE) to allow robust microbial or pathogen responses (Figure 3A) (54).

**Figure 3:**
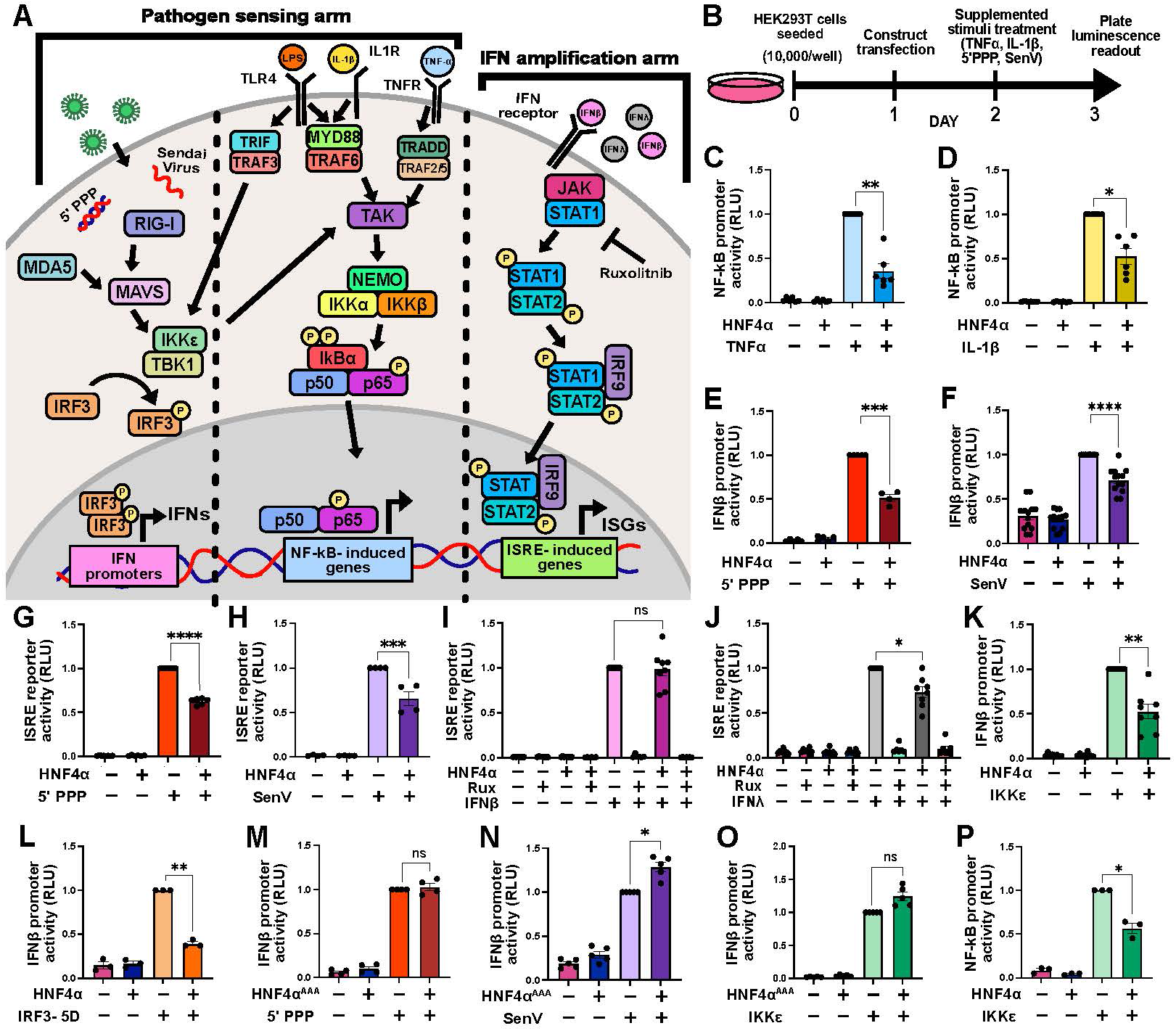
HNF4 suppresses IFN signaling in HEK293T cells. **(A)** Schematic of the IFN induction (pathogen sensing) and ISG induction (IFN amplification) pathways. **(B)** Schematic of the luciferase reporter assay. HEK293T cells were co-transfected with a firefly luciferase reporter driven by the indicated promoter, a Renilla normalization control, and either HNF4α or empty vector, then stimulated with an IFN inducing construct or supplemented stimulus. **(C-M)** Promoter activity in cells expressing HNF4α or empty vector after stimulation with the indicated stimuli. Activity is reported as relative light units (RLU), normalized to Renilla luciferase and to the stimulated treatment group. **(N-P)** IFNβ promoter activity in HEK293T cells expressing empty vector, wild-type HNF4α, or the DNA-binding domain mutant, HNF4αAAA, after stimulation with the indicated IFN-inducing stimuli. Each dot represents one independent experiment (mean of 3 technical replicates); bars show mean ± SEM. N = 3-13 independent experiments per panel. Statistics: repeated measures one-way ANOVA with Tukey’s multiple comparisons test. ns, P > 0.05; *P ≤ 0.05; **P ≤ 0.01; ***P ≤ 0.001; ****P ≤ 0.0001.

To test whether HNF4 regulates ISG expression, we overexpressed HNF4 in HEK293T cells, stimulated the NF-κB, IRF, and IFN pathways with known activators, and measured downstream promoter activity by luciferase assay (Figure 3B) (56–60). Because HEK293T cells express little or no endogenous HNF4 (61–63), we transfected them with either an HNF4 expression construct or an empty control construct, together with a luciferase reporter driven by the NF-κB or IFNβ promoter (Figure 3B). Twenty-four hours later, we stimulated the cells with an NF-κB activator (TNFα or IL-1β) or with a potent RIG-I activator that induces IFNβ (5′ PPP or Sendai virus [SenV]), and after another twenty-four hours, measured luciferase activity by plate luminescence. The presence of HNF4 significantly attenuated stimulus induced activation of both the NF-κB and IFNβ promoter reporters (Figure 3C-F). We similarly examined luciferase activity driven by the ISRE, a readout of the IFN amplification arm (54). HNF4α overexpression also led to a significant reduction in ISRE reporter-driven luciferase activity following stimulation with 5′PPP and SenV (Figure 3G-H), though IL-1β and TNF failed to induce robust ISRE activity, as expected (Supplemental Figure 5A,B). Together, these data show that HNF4 overexpression suppresses IFN and NF-κB induction *in vitro*.

### HNF4 suppresses ISG induction upstream of IFN production

HNF4 suppressed both IFNβ and ISRE activity, so we next asked where it acts in the pathways leading to ISG induction (Figure 3A). We transfected HEK293T cells with an ISRE-driven luciferase reporter, with or without HNF4. Twenty-four hours later, we treated the cells with IFNβ or IFNλ, which stimulates only the IFN amplification arm, and the following day we measured luciferase activity by plate luminescence as before. HNF4 did not suppress ISRE activity in the presence of IFNβ, but it partially suppressed IFNλ-induced activity (Figure 3I-J). To confirm that IFN signaling acted through the canonical JAK/STAT pathway, we also treated cells with ruxolitinib, a JAK inhibitor (64). As expected, ruxolitinib abolished ISRE reporter activity (Figure 3I-J). These data establish that HNF4 suppresses ISRE activity upstream of IFNβ and IFNλ induction, with a smaller additional effect on JAK/STAT signaling downstream of IFNλ *in vitro*.

### HNF4 DBD is required to suppress IFNβ and NF-κB activity

To determine where in the pathogen sensing arm HNF4 was acting, we transfected cells with the IFNβ promoter-driven luciferase and IKKε or a constitutively active IRF3 (IRF3-5D), both critical mediators of the IFNβ induction pathway, in the presence or absence of HNF4 (65). HNF4 was able to suppress IFNβ promoter and ISRE reporter activity downstream of both IKKε and IRF3-5D, suggesting HNF4 ’s suppressive activity occurred in the nucleus or by sequestering activated IRF3 outside of the nucleus (Figure 3K-L + Supplemental Figure 5C,D). Validating these results, and supporting the relevance of HNF4 suppression for type III as well as type I interferon, HNF4 suppressed IKKε-induced expression of *IFNB1*, *IFNL1*, and *ISG15* by qRT-PCR (Supplemental Figure 5E-G). We next examined the subcellular localization of HNF4 in the presence or absence of IKKε and found consistent nuclear localization of HNF4 localization across conditions, supporting the notion that HNF4 ’s suppression of IFNβ activity is likely to occur in the nucleus (Supplemental Figure 5H-K).

Because HNF4 is a well-characterized epithelial nuclear receptor and transcription factor, we hypothesized that it suppresses IFN induction through its DNA-binding domain (DBD) and transcriptional regulation. To test this, we generated a mutant HNF4 construct with three continuous point-mutations in the DBD of HNF4 (D105A, K106A, D107A) (66, 67). Alphafold structure prediction and subsequent ChimeraX analysis identified that the wild-type (WT) and mutated proteins have an angstrom RMSD score of 0.879 in defined regions, suggesting they are nearly identical in structure (Supplemental Figure 6A). Similarly, the HNF4 mutated construct was recognized by HNF4 antibodies by western blot and immunofluorescence and localized to the nucleus (Supplemental Figure 6B,C). Taken together, these data indicate that the HNF4 DBD domain mutant construct (HNF4 ^AAA^) does not significantly alter protein structure. We next sought to validate that our HNF4 ^AAA^ construct no longer induced HNF4 target genes. We transfected HEK293T cells with a luciferase reporter driven by the promoter of the HNF4 -induced gene, *TREH*, together with either the WT HNF4 or the HNF4 ^AAA^ construct (68). As expected, WT HNF4A strongly induced *TREH* promoter activity while HNF4 ^AAA^ induced significantly less activity (Supplemental Figure 6D). We next tested whether HNF4 ^AAA^ suppressed IFNβ promoter activity as effectively as WT HNF4 and found HNF4 ^AAA^ was no longer able to suppress IFNβ induction after stimulation with SenV, 5’PPP, or IKKε (Figure 3M-O). Similar results were obtained using the ISRE reporter (Supplemental Figure 6E-G). These results demonstrate that HNF4 needs to bind DNA to suppress IFN signaling.

We have demonstrated that HNF4 was able to suppress NF-κB-driven luciferase activity downstream of MYD88 (IL1β stimulation) and TRADD signaling (TNFα stimulation; Figure 3C,D). Though primarily known to induce IFN signaling, IKKε has been shown to co-stimulate the NF-κB signaling pathway downstream of MYD88 and TRADD by activating TAK in myeloid cells (Figure 3A) (52). We were therefore curious if IKKε could similarly induce NF-κB signaling in HEK293T cells. We found that IKKε was not only able to induce NF-κB promoter-driven luciferase activity, but HNF4 was able to significantly suppress this induction (Figure 3P). This is consistent with HNF4 acting at or downstream of IKKε-induced TAK signaling in the NF-κB pathway. To demonstrate IKKε-induced NF-κB promoter activity was likely occurring through true NF-κB signaling and not unmapped IFN signaling, we repeated the experiment with IRF3-5D and found that, as expected, constitutively active IRF3 did not induce NF-κB promoter activity (Supplemental Figure 6H). As HNF4 required a functional DBD to successfully suppress IFNβ-induced activity, we tested if the DBD was also required for NF-κB signaling suppression using our HNF4 ^AAA^ construct. Normalized plate luminescence readings showed that, like IFNβ-driven activity, HNF4 ^AAA^ was no longer able to suppress NF-κB promoter activity after stimulation with IL1β, TNF, or IKKε (Supplemental Figure 6I-K). Together, these data support that HNF4 can suppress IFNβ and NF-κB induction through its DBD domain *in vitro*.

### HNF4 induces candidate secondary mediators *in vitro*

HNF4 is known to act predominantly as an activator of transcription for epithelial genes in the intestine, kidney, and liver, but it is not known to bind extensively at many of the microbially-induced gene loci (20, 22, 35, 69). We searched our IFNβ- and NF-κB-driven luciferase constructs for putative HNF4 binding sites and found no putative HNF4 TFBS motifs in the promoter sequences of these constructs or HNF4a ChIP-seq peaks at these loci in mouse IECs (not shown) (24). This suggests HNF4 does not act directly on the IFNβ or NF-κB reporters, but instead regulates expression of secondary mediators that suppress IFNβ and NF-κB activity. We therefore sought to identify HNF4 -induced genes that might act as secondary, more direct, suppressors of inflammatory signaling. HEK293T cells were transfected with HNF4 and either an empty vector or IKKε, which induces activity across all reporter constructs (Figure 3K,P + Supplemental Figure 5C + Figure 4A). At 48 hours post-transfection, RNA was extracted and subjected to bulk RNA sequencing, and differential expression was compared between conditions (Supplemental Table 5). In Principle Component Analysis (PCA), control, HNF4, and HNF4 + IKKε samples clustered together, separate from IKKε samples (Figure 4B). Gene expression data suggest that HNF4 co-expression returns IKKε-stimulated samples to a more baseline-like state (Figure 4B-E + Supplemental Figure 7A,B). HNF4 and IKKε each upregulated their known target genes and expression changes were similar between replicates (Figure 4E). Consistent with luciferase results, HNF4 co-expression attenuated the expression of most ISGs induced by IKKε (Supplemental Figure 5C,E-G + Figure 4D-G). GO Term type analysis identified inflammatory response pathways among genes that were upregulated by IKKε and subsequently downregulated by HNF4 + IKKε, including RIG-I-like receptor signaling, positive regulation of canonical NF-κB signaling, and regulation of pattern-recognition receptor signaling (Figure 4H). Further supporting both the validity of this model and the established roles of HNF4, HNF4 overexpression downregulated pathways associated with cell division and cellular activation (24, 35). Co-expression of HNF4 + IKKε suppressed nearly all ISGs, with the notable exception of *ISG20*, whose expression remained unchanged, suggesting HNF4 activity was not immediately driving broad cell type transcriptional changes, but operating with specificity for IKKε-induced genes (Figure 4E). Interestingly, co-expression of HNF4 + IKK-ε increased *HNF4A* RNA levels, a finding that was confirmed at the protein level by western blot analysis (Figure 4E-G + Supplemental Figure 7C,D). Finally, sequencing analysis identified 18 genes significantly upregulated by HNF4 (HNF4 or HNF4 + IKKε co-expression) that might function as secondary mediators of IKKε suppression (“HNF4 induced” or “HNF4 + IKKε induced” in Figure 4E).

**Figure 4:**
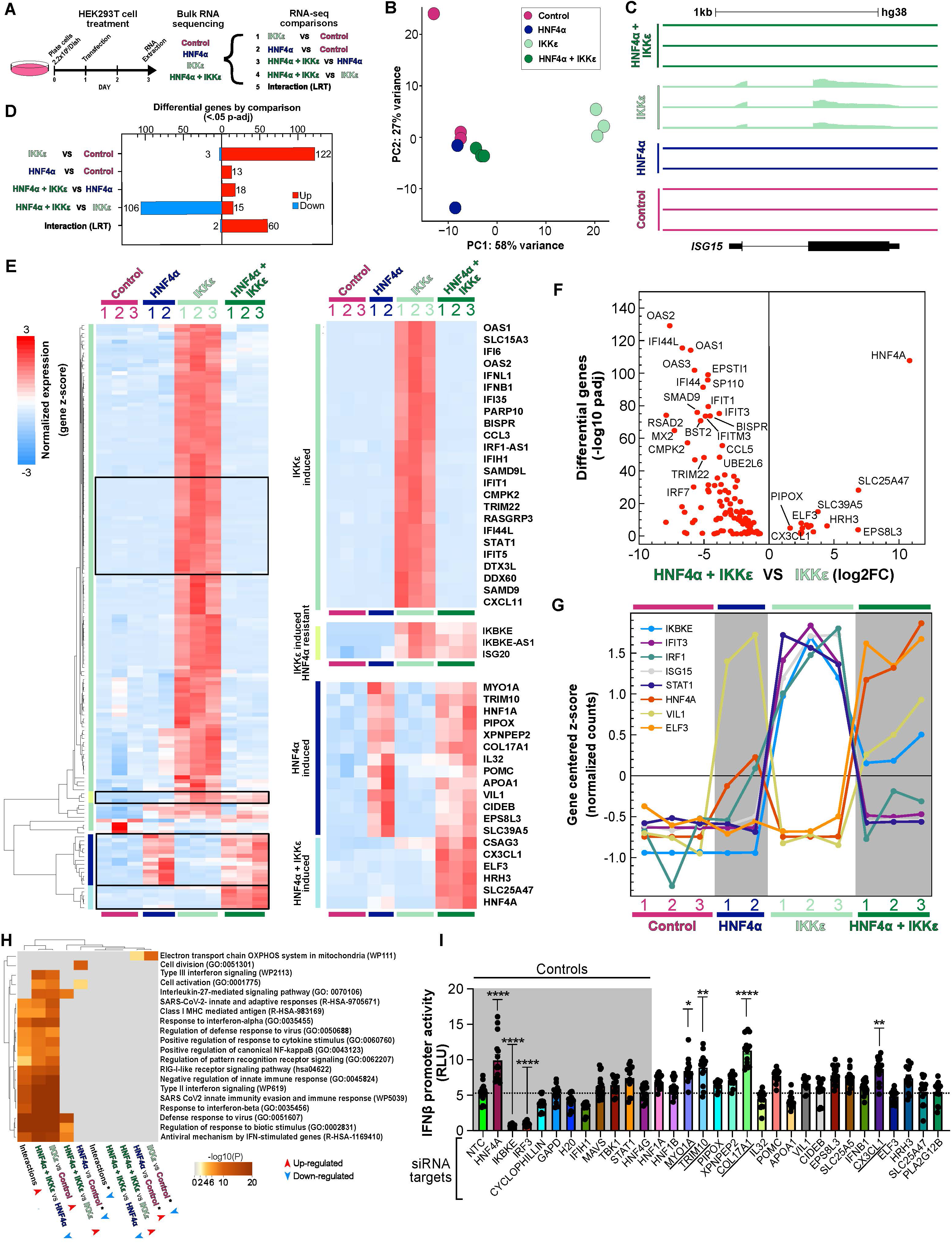
HNF4 induces candidate secondary mediators of IFN suppression. **(A)** Experimental design for bulk RNA-seq. HEK293T cells were transfected with an empty vector (control), HNF4α, IKKε, or HNF4α + IKKε, and RNA was collected 48 post-transfection. **(B)** Principal component analysis (PCA) of bulk RNA-seq samples. **(C)** Genome browser view of RNA-seq read coverage at the *ISG15* locus (hg38). Tracks show individual replicates and are scaled identically. **(D)** Number of significantly differentially expressed genes (DEGs; adjusted P < 0.05) for each indicated comparison, split into up-regulated (red) and down-regulated (blue). The interaction term was tested with a likelihood ratio test (LRT). **(E)** Heatmap of row-scaled expression (z-score of normalized counts) for the DEGs in (D). Expanded views show gene clusters that are IKKε induced, IKKε induced and HNF4α resistant, HNF4α induced, and HNF4α + IKKε induced. **(F)** Volcano plot of DEGs between HNF4α + IKKε and IKKε samples. **(G)** Z-scored normalized counts across samples for selected ISGs and HNF4α targets. **(H)** Enriched pathways among up- and down-regulated genes for each comparison in (D). Due to low numbers of significant genes in certain directional comparisons, reduced thresholds were used to capture potential biology (DEG thresholds; < .01 p-value >15 base mean). Color indicates −log10(P); arrowheads indicate the direction of regulation. **(I)** IFNβ promoter activity in HEK293T cells expressing HNF4α and IKKε after pretreatment with siRNAs targeting HNF4α-induced genes. Activity is reported as relative light units (RLU), normalized to Renilla luciferase. Gray shading marks control siRNAs [NTC, *HNF4A*, *IKBKE*, *IRF3*, etc]; the dashed line marks the NTC mean. Each dot represents one experimental replicate, bars show mean ± SEM; N = 5 independent experiments. Statistics for (I): one-way ANOVA with Dunnett’s multiple comparisons test versus NTC. *P ≤ 0.05; **P ≤ 0.01; ***P ≤ 0.001; ****P ≤ 0.0001.

To assess the functional relevance of each candidate, we obtained siRNAs targeting each of the 18 genes, plus additional controls, and tested whether their depletion impaired HNF4 induced suppression of inflammatory responses. Under HNF4 + IKKε co-expression conditions, we knocked down each candidate gene individually and measured IFNβ promoter-driven luciferase activity. Reference controls behaved as anticipated: knockdown of *HNF4A* increased IFNβ promoter activity, whereas knockdown of *IKBKE* or *IRF3* decreased activity (Figure 4I). Of all 18 candidate genes, only knockdown of *COL17A1*, *CX3CL1*, *MYO1A*, and *TRIM10* significantly impaired HNF4 -mediated suppression of IFNβ promoter activity when aggregated across 5 independent experimental replicates (Figure 4I). Taken together, these findings support that HNF4 induces expression of secondary mediators (*COL17A1*, *CX3CL1*, *MYO1A*, *TRIM10*) required for its suppression of IFN activity *in vitro*.

### Co-expression of HNF4 induced genes suppresses inflammatory signaling *in vitro*

Our siRNA screen demonstrated that HNF4 induced genes *COL17A1*, *CX3CL1*, *MYO1A*, and *TRIM10* are required for HNF4 -mediated IFN suppression. We next asked whether, in the absence of HNF4, overexpression of these gene hits was sufficient to suppress inflammatory signaling *in vitro*. For each of the four gene hits, we generated a construct overexpressing the respective cDNA from the constitutive CMV promoter. To ascertain their collective sufficiency to suppress inflammatory signaling, we co-expressed all four constructs (4 hits) and stimulated cells with different inflammatory stimuli (Figure 5A-E). As with HNF4, co-expression of the 4 hits was sufficient to suppress luciferase activity driven by IFNβ, NF-κB, and ISRE reporters in all stimuli tested (Figure 5A-E + Supplemental Figure 8A-C). To begin to determine if any of the 4 hits were individually sufficient to suppress inflammatory signaling, we expressed each hit alone and stimulated with IKKε. Surprisingly, no single hit suppressed IKKε-driven IFNβ or ISRE activity (Figure 5F-I + Supplemental Figure 8D-G). Similarly, co-expression of pairs of hits failed to suppress IKKε-induced signaling (Supplemental Figure 8H-S). These data indicate that IKKε-induced signaling of IFNβ-, NF-κB-, and ISRE-driven luciferase activity can only be suppressed by the co-expression of all 4 hits.

**Figure 5:**
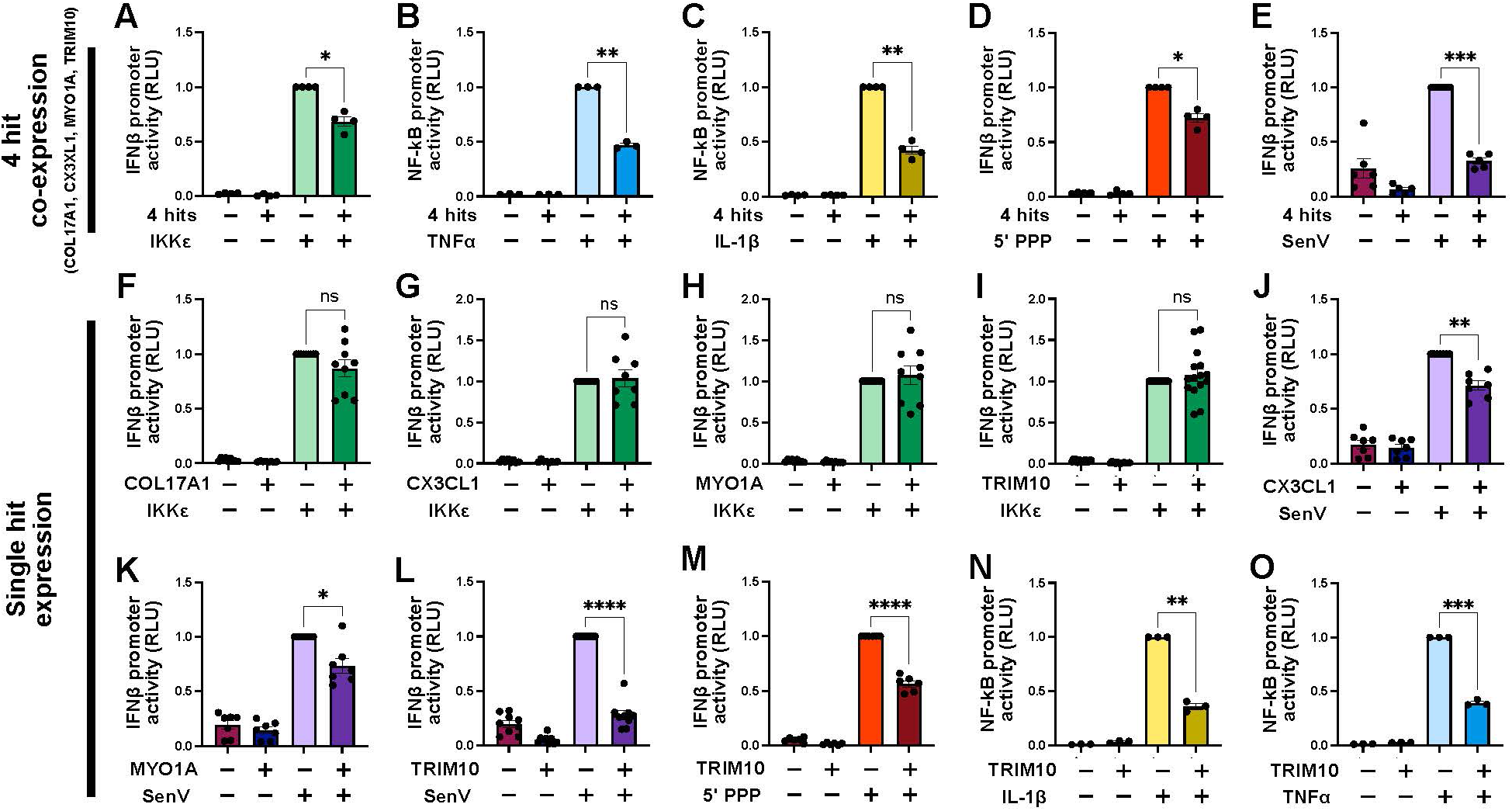
Co-expression of HNF4 -induced candidate secondary mediators suppress IFN signaling. **(A-E)** Promoter activity in HEK293T cells co-expressing genes targeted by the four siRNA screen hits (*COL17A1*, *CX3CL1*, *MYO1A*, and *TRIM10*; “4 Hits”) or empty vector, with or without the indicated stimulus. **(F-O)** Promoter activity in cells expressing individual hits or an empty vector control, with or without the indicated stimulus. Cells were co-transfected with a firefly luciferase reporter and a Renilla control and stimulated for 24hr (SenV, 5’PPP, TNF, IL-1β) or 48hr (IKKε). Activity is reported as relative light units (RLU), normalized to Renilla luciferase and to the stimulated treatment group. Each dot represents one independent experiment (mean of 3 technical replicates); bars show mean ± SEM. N = 3-13 independent experiments per panel. Statistics: repeated measures one-way ANOVA with Tukey’s multiple comparisons test. ns, P > 0.05; *P ≤ 0.05; **P ≤ 0.01; ***P ≤ 0.001; ****P ≤ 0.0001.

We next sought to determine if inflammatory signaling upstream of IKKε could be suppressed by individual hits. When cells were stimulated with SenV, only individual overexpression of *CX3CL1*, *MYO1A*, or *TRIM10* was sufficient to suppress IFNβ-promoter driven luciferase activity (Figure 5J-L + Supplemental Figure 8T). Under the same conditions, however, suppression of ISRE-promoter driven luciferase activity was observed only with *CX3CL1* and *TRIM10* overexpression (Supplemental Figure 8U-X). In contrast, treatment with 5′PPP revealed significant suppression of both IFNβ- and ISRE-promoter activity only after *TRIM10* overexpression (Figure 5M + Supplemental Figure 8Y-Ae). Similarly, *TRIM10* was the only gene whose overexpression was sufficient to suppress IL1β or TNF induced NF-κB activity (Figure 5N-O + Supplemental Figure 8Af-Ak). Together these results support that HNF4 induced suppression of IFNβ and NF-κB signaling is mediated by the combined action of multiple, downstream HNF4 target genes (*COL17A1*, *CX3CL1*, *MYO1A*, *TRIM10*). Additionally, whereas suppression of IFNβ- and NF-κB signaling upstream of IKKε was achieved to varying degrees by expression of individual HNF4 target genes, suppression of IKKε-induced inflammatory signaling required the co-expression of all 4 genes.

### HNF4 deletion alters ISG expression and pathogen response in intestinal epithelial cells

Cumulatively, our HEK293T cell culture studies established that HNF4 indirectly suppresses IFN and other inflammatory signaling pathways within the same cell by driving expression of multiple secondary mediators. However, the relevance of these interactions *in*

*vivo* remained unknown.

We therefore sought to determine whether HNF4 similarly suppresses IFN signaling in the intestinal epithelium. To do so, we generated jejunal organoids from SPF mice lacking HNF4 in their intestinal epithelium (HNF4 KO; *Vil:Cre*^+/-^; *Hnf4a*^F/F^) or their WT littermate controls (*Vil:Cre*^-/-^; *Hnf4a*^F/F^; Figure 6A). Quantitative RT-PCR revealed that organoids from HNF4 KO animals exhibited significantly higher baseline *ISG15* expression than WT animals (Figure 6B). Mirroring our cell culture data, HNF4 KO organoids maintained higher *ISG15* expression profiles after stimulation with viral mimetic polyinosinic:polycytidylic acid (poly I:C), but not IFNβ or IFNλ (Figure 6B). These data imply that, similar to HEK293T cells *in vitro*, HNF4 may also play a role in suppressing the pathogen sensing arm of IFN signaling at baseline in IECs.

**Figure 6:**
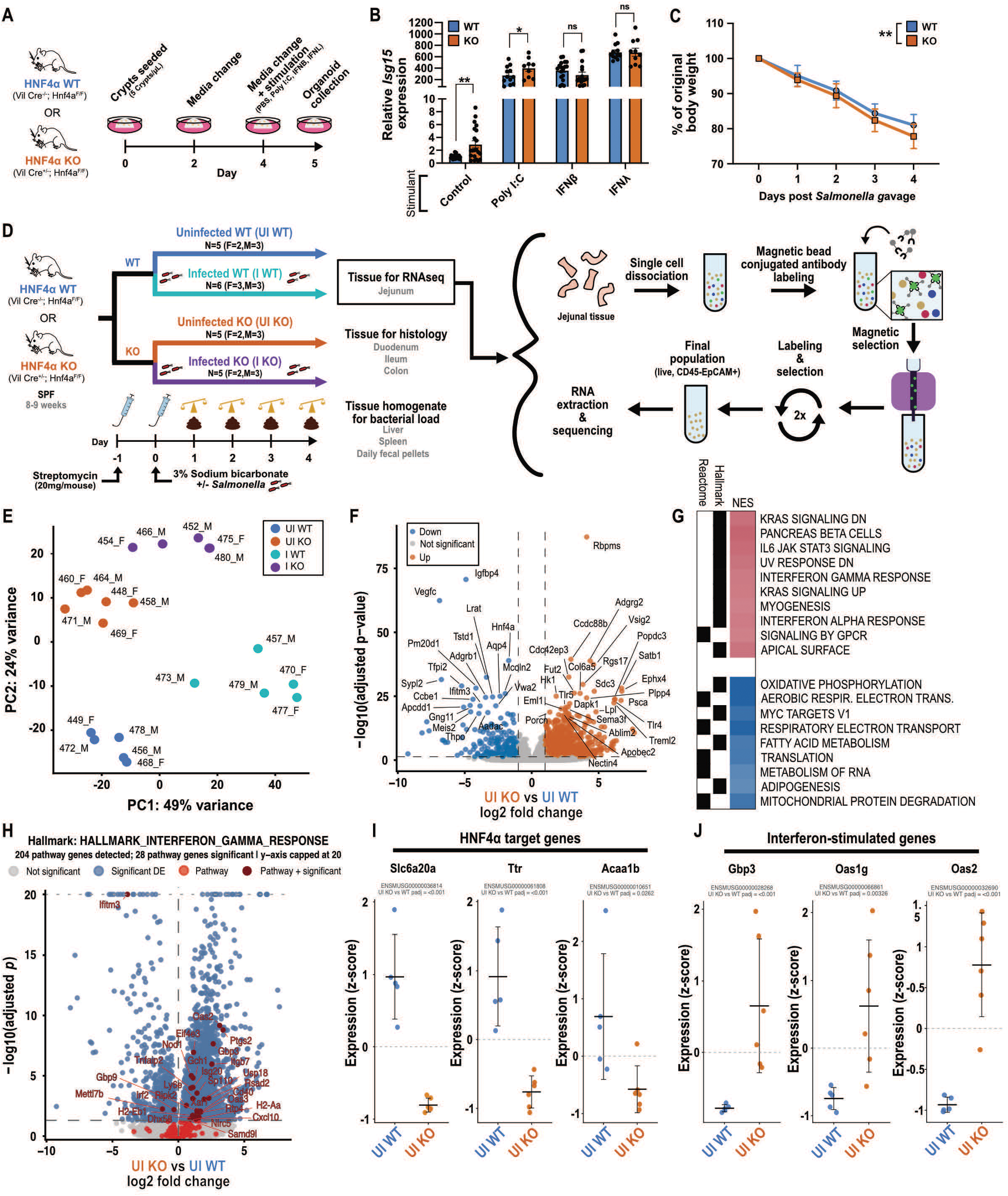
HNF4α KO IECs have elevated baseline ISG expression compared to WT controls. **(A)** Organoid experimental design. Jejunal crypts from SPF HNF4α KO or WT mice were seeded (5 crypts/µL) on day 0, with media changes on days 2 and 4. On day 4, organoids were stimulated with PBS (control), poly I:C, IFNβ, or IFNλ, and RNA was collected 24 h later for qRT-PCR. **(B)** *Isg15* expression in WT and KO organoids after the indicated stimulation, measured by qRT-PCR relative to *Gapdh* and normalized to unstimulated WT. Each dot represents one well (n = 12-21 wells per group) from organoid lines derived from 4-7 animals; bars show mean ± SEM. Statistics: paired two-tailed t-test between matched genotype controls. **(C)** Body weights after *Salmonella enterica* serovar Typhimurium (*S.* Tm) gavage, shown as percent of starting weight, in SPF WT and KO mice. Symbols show the group mean ± SEM (n = 23-25 animals per group, pooled from 3 independent experiments). Statistics: two-way repeated measures ANOVA with Šídák’s multiple comparisons test between WT vs. KO at each time point. **(D)** *S.* Tm infection design and IEC isolation. Group sizes: uninfected (UI) WT n = 5 (2F, 3M), infected (I) WT n = 6 (3F, 3M), UI KO n = 5 (2F, 3M), I KO n = 5 (2F, 3M). **(E)** PCA of IEC RNA-seq samples. **(F)** DEGs between UI KO and UI WT IECs. **(G)** Gene set enrichment analysis (GSEA) of UI KO vs. UI WT IECs. **(H)** Volcano plot from (F) with Hallmark Interferon Gamma Response genes highlighted. The y-axis is capped at 20. **(I-J)** Z-scored normalized expression of selected HNF4α target genes (I) and ISGs (J) in UI WT and UI KO IECs. ns, P > 0.05; *P ≤ 0.05; **P ≤ 0.01; ***P ≤ 0.001; ****P ≤ 0.0001.

Given the elevated baseline ISG expression in HNF4 KO organoids and the importance of IFN signaling and ISG expression in pathogen response, we next asked whether HNF4 deletion affects the IFN response *in vivo* during homeostasis and infection. As HNF4 was able to suppress the IFN response in both viral and bacterial-induced pathways *in vitro*, we infected HNF4 KO animals with different IFN-stimulating pathogen models (14, 70, 71). HNF4 KO and WT mice infected with Coxsackie B virus did not differ in weight loss or viral load at acute or chronic infection timepoints (Supplemental Figure 9K-M). In rotavirus infection, HNF4 KO mice similarly did not show differences in weight loss, but did have a significant increase in rotavirus genome copies one day earlier than WT controls (Supplemental Figure 9A-M). *Salmonella enterica* serovar Typhimurium (*S*. Tm.) infection caused a modest, but significant, increase in weight loss in HNF4 KO animals, accompanied by shortened jejunal villi and increased distal colon crypt depth (Figure 6C + Supplemental Figure 9N-V). As weight loss and villus blunting are associated with inflammatory states, we proceeded to examine how HNF4 loss impacts IFN signaling and other transcriptional responses to *S*. Tm. infection (72–74).

We isolated IECs from uninfected (UI) and *S*. Tm. infected (I) HNF4 KO and WT animals, enriched for live, CD45^-^, EpCAM^+^ cells with magnetic sorting, and performed bulk RNA-sequencing (Figure 6D + Supplemental Figure 10A-I). PCA analysis showed samples clustered distinctly from one another, implying substantial transcriptional impact from both infection and HNF4 status (Figure 6E). We first sought to examine gene expression changes between uninfected HNF4 KO and WT animals (KO/WT (UI)). Compared to WT, GSEA analysis showed HNF4 KO IECs were enriched in genes associated with IFN response and JAK/STAT3 signaling (Figure 6F-J). As expected, downregulated genes included HNF4, HNF4 target genes, and genes associated with metabolism (Figure 6F,G,I) (24, 35). In accord with our cell culture and organoid results, these data indicate that HNF4 KO IECs have higher expression of inflammatory programs, including ISG expression, at homeostatic baseline.

We next expanded comparisons amongst the four conditions (UIWT, UIKO, IWT, IKO) to interrogate the impact of HNF4 loss on the response to *S*. Tm. infection in IECs. Direct comparison of the loss of HNF4 to WT with and without infection [KO/WT(UI) vs. KO/WT(I)] identified a grossly analogous transcriptional response (Figure 7A + Supplementary Table 6). Similarly, the response of *S*. Tm. infection in WT and KO [I/UI (WT) vs. I/UI (KO)] was generally correlated and all infected animals showed induction of many *Reg3* family genes regardless of genotype (Figure 7B-C). However, when we performed interaction analysis across all four conditions, we identified numerous genes suggesting an impaired response to infection in HNF4 KO animals [KO/WT (I)] (Figure 7A-D). GSEA analysis identified that a large group of ISGs failed to be completely induced in IKO, including *Irf7*, *Isg15*, and *Oas3*. This suggests a fundamental reduction in IFN-based responses in infected KO relative to infected WT conditions despite consistent, modest activation of ISGs in UIKO relative to UIWT (Figure 7D-G + Supplementary Table 6). Interestingly, several non-ISG genes (*Bmp2, Bmp3, Pla2g5, Snx7*) also showed significant interactions highlighting potential programs or regulators that were also compromised (Figure 7F-G).

**Figure 7:**
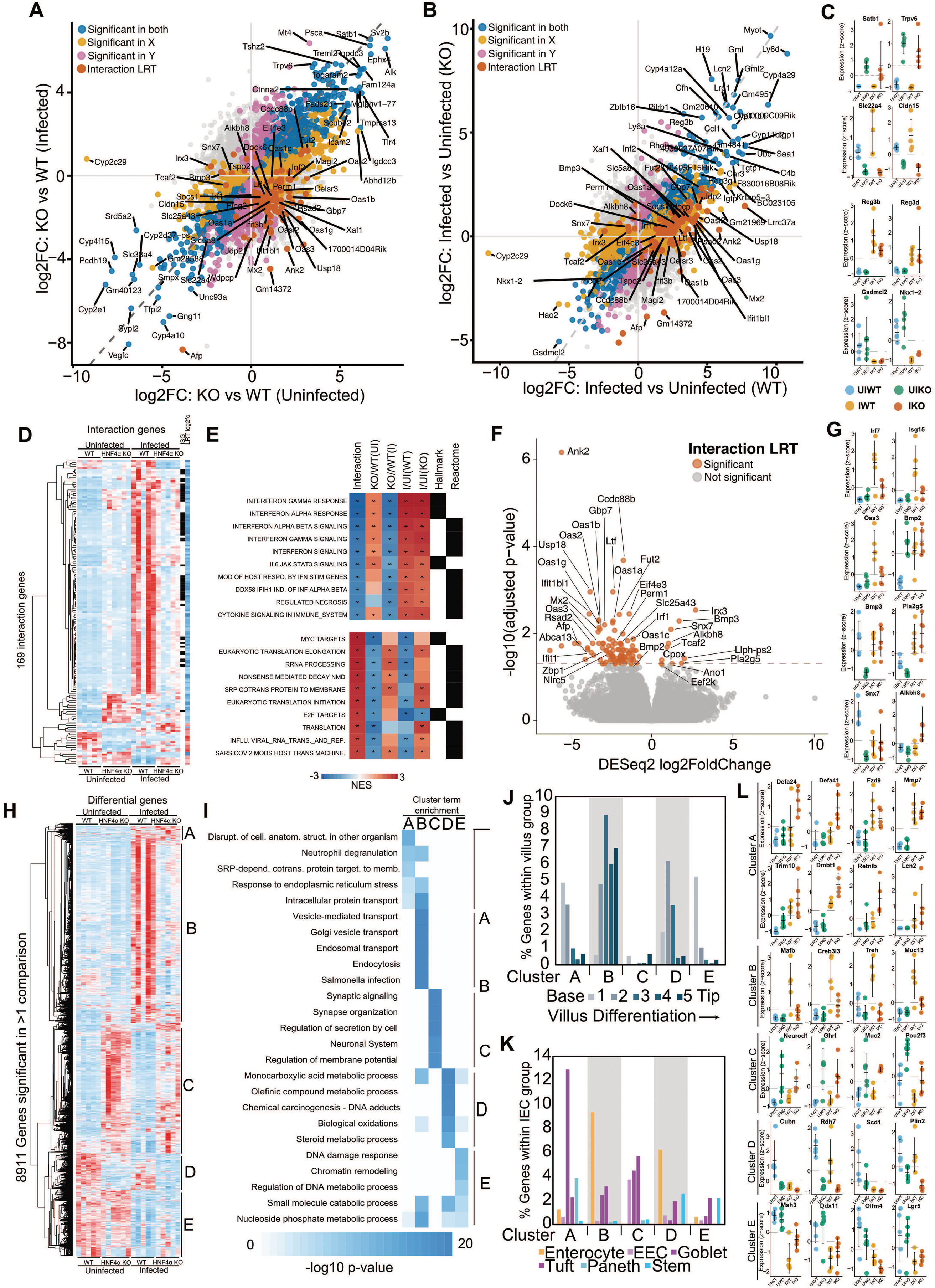
Loss of HNF4α limits induction of enterocyte ISGs and increases secretory IEC defenses following *S.* Tm. **(A)** Scatterplot comparing the expression log2FC effect in IECs of HNF4α loss to WT in uninfected (KO/WT (UI)) and *S.* Tm. infected (KO/WT (I)) SPF mice. **(B)** Scatterplot comparing the expression log2FC effect in IECs of *S.* Tm. infected mice to uninfected mice in WT (I/UI (WT)) and HNF4α KO (I/UI (KO)) mice. **(C)** Plot of z-scored expression of IEC normalized counts from example genes that are up or down regulated by HNF4α KO or infection, without apparent interaction between HNF4α genotype and *S.* Tm. infection status. **(D)** Hierarchically clustered heatmap of row z-scored normalized counts for genes that are significant < .05 p-adj for LRT or Wald interaction tests. Genes are marked for membership in Hallmark ISG and significant induction by interferon lambda in small intestinal organoids. Interaction log2FC is also included for each gene. **(E)** Heatmap of GSEA Hallmark and Reactome output for the top 10 significant low NES or high NES terms based on genes ordered by the interaction analysis. GSEA for each of the four comparisons represented in (A) and (B) are included and significant GSEA results are marked with an asterisk. **(F)** Volcano plot of significant interaction genes (interaction log2FC). **(G)** Plot of z-scored expression of IEC normalized counts from example interactions genes. **(H)** Hierarchically clustered heatmap of row z-scored normalized counts for genes that are significant in at least one of the four comparisons represented in (A) and (B). Major clusters are labeled from A-E. **(I)** Heatmap of the top 5 Metascape terms for gene lists for each of the 5 subclusters from (H). Proportion of genes from each cluster that have membership in **(J)** villus differentiation group (5) or **(K)** IEC subtype(75). **(L)** Plot of z-scored expression of IEC normalized counts from genes from the 5 clusters in (H).

We broadened our analysis by clustering genes that were differential in any comparison and found numerous gene expression patterns that implied a unique response to *S*. Tm. (Figure 7H). Unlike ISGs, a subset of infection induced genes showed maintained or increased expression in the IKO condition (Cluster A). These included Paneth cell genes, *Fzd9* and *Mmp7*, and genes involved in direct antagonism of bacterial pathogens, like many defensins (e.g. *Defa24*, *Defa41*), *Dmbt1*, *Retnlb*, *Lcn2*, and *Trim10* (Figure 7I). These additional, seemingly transcriptionally separated from ISGs, defense responses may explain how HNF4 KO animals suppress *S*. Tm. pathogenesis when ISG induction is limited following infection. We found ISGs were part of a larger group of genes (Cluster B) that, while not all significant for interaction, showed failed maximal induction comparing IKO to IWT (Figure 7H). The level of relative induction of these genes in IWT dwarfs subtler changes in other conditions and comparisons. Importantly, groups of genes that were upregulated (Cluster C; enriched in secretion and synapse terms) or downregulated (Cluster D; enriched in metabolism terms) across KO animals or suppressed by infection (Cluster E; enriched in DNA damage) typically showed limited interaction between infection status and genotype (Figure 7H).

Based on HNF4 ’s role in enterocyte differentiation and identification of anecdotal secretory and Paneth cell genes, we queried these clusters for their membership along the villus differentiation axis (5) and separately within IEC subtype groups (Figure 7J-K) (75). In Cluster A, we identified interpretable patterns of secretory goblet and Paneth cell genes with limited villus differentiation that showed maintained or increased expression in IKO. In contrast, Cluster B genes that failed to be fully induced in IKO (e.g. *Mafb*, *Creb3l3*, *Treh*, *Apoa1*) suggests that *S*. Tm. infection promotes villus enterocyte differentiation. However, HNF4 may be required for full induction of ISGs and shifting to a more mature enterocyte-heavy population during *S*. Tm. infection (Figure 7J-L). Consistent with this, Cluster C genes which are upregulated regardless of infection in HNF4 KO mice, were not enriched in enterocyte genes and instead included EEC, goblet, and tuft associated genes (e.g. *Neurod1*, *Ghrl*, *Muc2*, *Pou2f3*). Cluster D, genes universally suppressed by infection, were enterocyte genes associated with expression closer to the villus base (e.g. *Cubn*, *Rdh7*, *Scd1*, *Plin2*) (5). Cluster E, which were consistently downregulated by infection, included DNA-damage genes *Msh3* and *Ddx11* and stem cell genes (e.g. *Lgr5* and *Olfm4*) (Figure 7L). This is consistent with a model where HNF4 loss reduces differentiated enterocyte programs and induces secretory lineage genes, both at baseline and after infection. Consistent with this model, though HNF4 loss frequently leads to induction of ISGs at baseline, during infection HNF4 KO animals cannot properly further upregulate the programs of terminal IEC differentiation and ISG induction which appear to be linked in the response to *Salmonella*.

## Discussion

Here, we sought to uncover how cells balance identity and function in the face of microbial challenges. We first asked how jejunal IECs respond transcriptionally, at single-cell resolution, to the diverse stimuli that accompany microbiota colonization and investigated a predominant mechanism behind this response. We found that after microbial stimulation, individual IECs broadly upregulate inflammatory programs (enriched for STAT, IRFs, NF-κB TFBS motifs) that are inversely correlated with metabolism-associated programs (enriched for HNF4 TFBS motifs). Supporting this, HNF4 deletion models often show upregulation of microbially induced genes and inflammatory programs, suggesting HNF4 may interact with these programs. We show that HNF4 suppresses the IFN response and ISG expression using the highly manipulable HEK293T cell platform, and that this novel pathway interaction likely also exists in the intestinal epithelium using mouse organoids. Finally, we provide evidence that disrupting this interaction through HNF4 deletion alters the IEC transcriptional landscape at baseline and after *Salmonella* infection. Together these results provide strong evidence for a novel HNF4 -IFN axis mediating communication between the distinct gene regulatory networks governing intestinal epithelial identity and microbial sensitivity.

Our scRNA-seq analysis reveal that microbial colonization induces a broadly shared transcriptional response across jejunal enterocytes rather than a program confined to discrete enterocyte subtypes. However, certain genes with high subtype expression specificity may still perform more specific functions in practice. For example, *Tm4sf4* is highly expressed in Clusters 1 and 5 despite it being significantly downregulated by microbes in all enterocyte clusters. The microbiota induced program was dominated by IFN-related and other inflammatory genes, matching earlier bulk studies of the microbial response (24, 35, 36). Cluster 6, which increased in relative proportion in two of three of our CV samples, expressed this program most strongly, but nearly every other enterocyte cluster upregulated it in the same direction. This suggests that microbiota colonization may induce increases in certain IEC cell types but can also induce inflammatory transcriptional programs that compete with programs driving IEC subtype identity and enterocyte differentiation.

Recent studies identified focal populations of ISG-high enterocytes located sporadically throughout the intestinal epithelium (12, 76). Cluster 6 resembles these cells in expression pattern, and we speculate that Cluster 6 is derived from these focal ISG-high villi in our dataset (12, 76). Prior work proposed that microbes stimulate immune cells to produce IFNλ, which then signals to epithelial cells and induces ISG-high states (12). However, our scRNA-seq data identified Cluster 6 cells in the GF jejunum, though at lower levels than in CV mice, contradicting a strictly microbe-dependent origin. Consistent with this, other works have also identified GF IECs expressing ISGs (77, 78). Our data also suggest that ISG expression is not driven by observable viral infection in GF animals. However, technical caveats such as sequencing depth limits and the absence of polyA tails in viruses such as flaviviruses, preclude conclusive evidence of this argument (79). Differences in ISG expression levels or in expression patterns of specific ISGs between GF and colonized animals may also account for the lack of ISG-high IEC populations at the macroscopic level (78, 80). How inflammatory-program induction occurs in the GF epithelium and if different microbes induce these programs variably remains an open question.

Our *in vitro* data suggest that HNF4 acts as a brake on inflammatory signaling in epithelial cells. This suppression appears to be mediated by the upregulation of multiple genes rather than a single effector. The 4 hits from our siRNA screen in HEK293T cells (COL17A1, CX3CL1, MYO1A, and TRIM10) have disparate functions, and no mechanistic connection among them is currently known. COL17A1 encodes Collagen XVII, a protein important for cell adhesion and motility (81, 82). MYO1A, a previously established HNF4 target, maintains brush border polarization and function in epithelial cells (20, 83). CX3CL1 is a well-known chemoattractant that recruits immune cells to sites of inflammation but also regulates cell-cell adhesion and can act as a viral mimetic (84, 85). TRIM10 belongs to the tripartite motif (TRIM) superfamily of E3 ligases, which ubiquitinate proteins and thereby target them for degradation (86). Given these varied functions, it is unlikely that the four hits act within a single mechanistic pathway to regulate inflammatory signaling. We propose instead that their individual functions cumulatively give HEK293T cells the functional capacity to suppress inflammatory signaling. This suggests a larger working model wherein the HNF4 regulon includes not only genes promoting the development and physiology of epithelia in the intestine and other organs, but also a battery of genes that interactively restrain IFN and other inflammatory pathways.

HNF4 is a known inducer of epithelial differentiation and influences both the proliferative and morphological state of HEK293T cells (20, 61). Because 3 of our 4 hits (COL17A1, CX3CL1, and MYO1A) support cell structure and adhesion, their roles in regulating inflammatory signaling may depend on maintaining a differentiated state that permits proper membrane protein presentation and cell motility and adhesion, rather than on direct participation in a signaling pathway. However, CX3CL1 may also act through a second mechanism as its tethered form contains a sialylated mucin-like stalk that can engage glycan-binding viruses (20, 87). This function is particularly relevant to our SenV findings, in which CX3CL1 overexpression alone suppressed SenV-induced IFNβ promoter activity. As Sendai virus adheres to epithelial membranes through sialylated glycans, CX3CL1’s stalk may be acting as a decoy that limits pathogen entry (88, 89).

Of the 4 hits, TRIM10 is the most likely to act directly in HNF4 -induced suppression of the IFN response. TRIM family members have long been associated with host defense and viral pathogenesis because their ubiquitination of target proteins can either enhance or suppress IFN and NF-κB signaling in a context-dependent manner (90, 91). Consistent with this, TRIM10 has been associated with both upregulation and downregulation of IFN signaling (92, 93). How TRIM10 interacts with the pathogen-sensing arm of our model to suppress IFN signaling, however, remains unclear.

Our organoid results further support a role for HNF4 in suppressing IFN signaling in the intestinal epithelium. Because HNF4 is required for stem cell renewal and maintenance in intestinal organoid culture (94, 95), we examined ISG expression only in primary jejunal organoid cultures supplemented with Wnt3a (94). In these cultures, loss of HNF4 significantly increased baseline ISG expression relative to WT organoids. This baseline increase was not fully penetrant, however as ISG expression was not always higher in every animal, consistent with variability seen in HNF4 KO animals (22, 27, 94–97). The significant overall difference nonetheless reinforces the concept that HNF4 suppresses inflammatory signaling at baseline, while the incomplete penetrance indicates that loss of HNF4 alone does not induce ISG expression. These data further suggest that the signals driving baseline ISG expression either persist transcriptionally in culture for at least six days or originate from the IECs themselves. Which factors are responsible for inducing variable ISG signatures and whether the HNF4 - induced mediators of suppression identified in HEK293T cells (COL17A1, CX3CL1, MYO1A, and TRIM10) play a similar role in organoid cultures remains to be determined.

Comparison of IECs from uninfected HNF4 KO and WT mice showed that loss of HNF4 is accompanied by higher baseline homeostatic ISG expression *in vivo*, extending our findings from cell lines and organoids to the intact intestinal epithelium. Loss of HNF4 also significantly upregulated BMP2 and downregulated BMP3. BMPs are known to guide differentiation in the intestine and have been tied to the control of inflammation (6, 98–102). In line with these changes, BMP2 signaling has been shown to promote inflammatory pathways, whereas BMP3 suppresses them (101, 103–105). Together, these results from uninfected mouse IECs further support the working model that HNF4 suppresses IFN responses and suggests a possible role for BMP signaling, though future studies will be needed to test this directly.

In response to infection with *S.* Tm., HNF4 KO animals mounted a dysregulated response relative to WT animals, successfully upregulating antibacterial genes but failing to upregulate ISGs. *Salmonella* infection is known to induce both epithelial differentiation and secretory cell density. Loss of either terminally differentiated cells expressing brush border proteins or secretory cells leads to worse infection outcomes (106–114). Because HNF4 is critical for the terminal differentiation of IECs and ISG-high cells are canonically found at the villus tip, we posit that ISG induction fails in HNF4 -knockout animals after *S.* Tm. infection because these animals can not sufficiently generate terminally differentiated IECs on these timescales (12, 21, 115). Consistent with this hypothesis, the genes that fail to be upregulated by *S.* Tm. infection in HNF4 KO animals are more highly expressed at the villus tip (5). Despite the failed upregulation of ISGs and reduced villus tip-associated genes, HNF4 KO animals did not show increased *S.* Tm. burden in fecal pellets or peripheral organs. We suspect that compensatory upregulation of antimicrobial and secretory cell-associated genes, such as defensins, *Trim10,* and *Muc2*, accounts for this outcome. Indeed, previous work has shown that HNF4 KO animals increase goblet cell numbers in response to non-invasive *Salmonella* infection (116). *Salmonella* infection results are supported by our observations in rotavirus infection. In rotavirus infection, the IFN response is believed to control initial viral shedding, while adaptive immunity resolves viral clearance (117). Though experimental confirmation is needed, it is therefore likely that the increased viral shedding observed in early infection in HNF4 KO animals is due to failed upregulation of the IFN response Understanding how HNF4 KO animals successfully compensate for the loss of IFN signaling in infection mechanistically will need careful investigation and may relate to specific pathogen dynamics. Potential roles for HNF4γ in that compensation and other HNF4 KO phenotypes also cannot be excluded, though we note that the HNF4 and HNF4γ have very similar patterns of DNA occupancy across the IEC genome (24).

*Trim10* expression was upregulated in HNF4 KO animals compared to WT animals regardless of infection status. While our HEK293T cell data supports that TRIM10 can be induced by HNF4, it can also be induced through IFNs (118, 119). As the SPF intestinal epithelium is known to have tonic IFN signaling, it is not surprising that TRIM10 is expressed, even in the absence of HNF4 (12). The pattern of TRIM10 expression in our data is consistent with antimicrobial proteins, suggesting it may be upregulated as a protective response to inflammation and infection. Further investigations will need to determine if TRIM10 acts to suppress IFN signaling *in vivo* and what regulates its transcription in IECs.

In summary, our findings support a model in which HNF4 -driven and inflammatory programs exist in tension within individual cells and interact to maintain intestinal homeostasis. Several questions remain open. Our mechanistic experiments relied on HNF4 overexpression in HEK293T cells, so whether COL17A1, CX3CL1, MYO1A, and TRIM10 mediate suppression in the intestinal epithelium is not yet known. The incomplete penetrance of ISG induction after HNF4 loss in mice further indicates that additional signals, such as microbes, determine whether loss of HNF4 results in inflammation. Defining these signals, and determining how differentiation state, BMP signaling, and secondary mediators each contribute to the observed HNF4 -IFN axis, will clarify how IECs balance nutrient absorption against pathogen defense. Understanding the balance and regulation of these transcriptional states may allow for development of new therapeutic targets aimed at altering IEC metabolism or inflammatory signaling.

HNF4α and IFN signaling have each been implicated in diverse gastrointestinal diseases, including inflammatory bowel disease (27, 28, 31–34, 120, 121), colorectal cancer (122, 123), celiac disease (124, 125), and viral infection (15, 126). The HNF4α-IFN interaction we describe here suggests that these two pathways may also intersect in those disease contexts, a possibility that future studies can test.

## Acknowledgments

We thank Dr. Mary Estes for technical advice for rotavirus experiments. The UNC-Chapel Hill CGIBD Advanced Analytics core contributed to our scRNA-seq experiments. Gabrielle Cannon prepared sorted cells and libraries for sequencing. We thank the Duke University School of Medicine for the use of the Sequencing and Genomic Technologies Shared Resource, which provided bulk RNA sequencing services for the HEK293T and *Salmonella* experiments. The Duke Cancer Institute Flow Cytometry Shared Resource provided assistance with flow cytometry analysis and experimental design related to the magnetic bead separation validation studies. We particularly acknowledge Dr. Bin Li for technical assistance with assay setup, controls, and data acquisition. We thank the Duke BioRepository and Precision Pathology Center Research Histology Laboratory for embedding, cutting, and staining histology slides. We thank the Duke Gnotobiotic Core and Duke Animal Care and Use program for their work in maintaining our germ-free and SPF colonies and help with colonization experiments. The Duke Light Microscopy Core Facility aided in the imaging studies. The TREH-promoter luciferase construct was a generous gift from Dr. Jesper Troelsen.

## Supplemental Figures

**Supplemental Figure 1: Gating strategy for FACs IEC isolation. (A)** Gating strategy and gate statistics **(B)** for jejunal IECs from GF and CV mice. **(C-G)** Gates C, M, and E-1 for unstained, single color, and FMO controls.

**Supplemental Figure 2: scRNA-seq of CV and GF mouse IECs. (A)** Each cell was assigned a module score for 5 published regional domains along the small intestine(127). The highest scoring regional domain represents each cell separately and is colored corresponding to the key. **(B)** As in (A) for 5 published regions in the villus differentiation program(5). Example genes that are significantly upregulated **(C)** or downregulated **(D)** in CV/GF in a limited number of clusters subset from heatmaps in Figure 1G. **(E)** Pseudobulk normalized counts from microbially responses genes highlighting potential relationships with enterocyte subtype-specific expression. **(F)** FeaturePlots for example genes in (E). **(G)** DimPlot by replicate to highlight cell cluster proportion. **(H)** FeaturePlot of GF and CV replicates for representative ISG, *Isg15*.

**Supplemental Figure 3 Quantification of transposable element expression. (A)** Differential expression of transposable element loci in cells from conventionally reared (CV) and germ-free (GF) mice using scRNA-seq data. **(B)** Enrichment of TE families among differentially expressed TE loci. Numbers indicate the number of differentially expressed loci from each family with log_2_FC ≥ 1 and adjusted p-values < 0.05. Asterisks indicate enriched TE families that are differentially expressed in CV and GF mice with FDR-adjusted p-values < 0.05.

**Supplemental Figure 4: A cluster 6 program correlates with microbially regulated genes at a cell by cell level. (A)** Cluster by cluster metrics split by GF and CV condition shows cluster 6 behaves within the normal range for median UMIs/cell (top), median features/cell (middle), and number of cells. **(B)** Scatterplot comparing scRNA-seq microbial response log2FC (CV/GF) in cluster 2 to cluster 2 find all markers log_2_FC colored by significance categories. **(C)** Scatterplot comparing scRNA-seq microbial response log2FC (CV/GF) in enterocytes to cluster 2 find all markers log_2_FC colored by significance categories. **(D-L)** Scatterplots highlighting variations on cell by cell correlations for each gene’s expression level in each cell to a cluster 6 module score of the top 50 significant cluster 6 find all markers genes ordered by log2FC compared to the gene’s microbial response in enterocytes (log_2_FC CV/GF). **(D)** All cells included, **(E)** correlation restricted to expression values from GF cells, **(F)** correlation restricted to expression values from CV cells, **(G)** correlation restricted to expression values from cluster 6 cells, **(H)** correlation restricted to expression values from non-cluster-6 cells, **(I)** example correlation between a maintained cluster 6 program and permutation of each gene’s expression values independently randomized across cells. **(J)** Correlation for all cells, removing the contribution of the top 50 cluster 6 marker genes to exclude the inherent correlation between these genes and the cluster 6 program, **(K)** correlation for each gene is restricted to cells with greater than 0 expression. **(L)** Correlation for genes using all cells as in (D) for the cluster 2 program compared to the gene’s microbial response in enterocytes (log_2_FC CV/GF). Scatterplot of HNF4α loss log2FC (HNF4α KO/WT) for two distinct genomewide intestinal expression datasets (47, 128) for **(M)** cluster 6 positive log2FC marker genes and **(N)** hallmark interferon response genes.

**Supplemental Figure 5: HNF4α suppresses ISG induction downstream of nuclear translocation in HEK293T cells. (A-D)** ISRE promoter activity in HEK293T cells expressing HNF4α or empty vector, with or without the indicated stimulus. Cells were co-transfected with a firefly luciferase reporter and a Renilla control and stimulated for 24 h (IL-1β, TNFα) or 48hr (IKKε, IRF3-5D). Activity is reported as relative light units (RLU), normalized to Renilla luciferase and to the stimulated treatment group. Each dot represents one independent experiment (mean of 3 technical replicates); bars show mean ± SEM. N = 3-13 independent experiments per panel. Statistics: repeated measures one-way ANOVA with Tukey’s multiple comparisons test. **(E-G)** *ISG15* (E), *IFNL1* (F), and *IFNB1* (G) expression measured by qRT-PCR in cells expressing HNF4α or empty vector, with or without IKKε. Expression is relative to *GAPDH* and normalized to HNF4α overexpression. Each dot represents one well (n = 3 wells per group); bars show mean ± SEM. Statistics: repeated measures one-way ANOVA with Tukey’s multiple comparisons test. **(H-K)** Immunofluorescence of HEK293T cells expressing IKKε alone (H), HNF4α alone (I), HNF4α + IKKε (J), or stained with IgG control antibodies (K). Cells were stained for HNF4α (green), IKKε (red), and nuclei (Hoechst, blue). Magenta boxes indicate regions shown at higher magnification. Images were acquired at 60×. Scale bars are represented on each image. ns, P > 0.05; *P ≤ 0.05; **P ≤ 0.01; ***P ≤ 0.001; ****P ≤ 0.0001.

**Supplemental Figure 6: Mutating the HNF4α DNA binding domain prevents suppression of IFN signaling in HEK293T cells. (A)** Predicted protein structure of WT HNF4α and the mutated HNF4α^AAA^constructs by ChimeraX. Green box represents inset for zoomed inset highlighting the DBD mutation. **(B)** Western blot staining of Vinculin and HNF4α in HEK293T cells expressing HNF4α, HNF4α^AAA^, or an empty vector control. **(C)** Immunofluorescence of HEK293T cells expressing HNF4α or HNF4α^AAA^. Cells were stained for HNF4α (green) and nuclei (Hoechst, blue). Images were acquired at 60×. Scale bars are represented on each image. **(D)** HNF4α target *TREH* promoter activity in HEK293T cells expressing HNF4α, HNF4α^AAA^, or an empty vector control. **(E-K)** Indicated promoter activity in cells expressing HNF4α^AAA^ or empty vector control, with or without indicated stimuli. Cells were co-transfected with a firefly luciferase reporter and a Renilla control and stimulated for 24hr (SenV, 5’PPP, TNF, IL-1β) or 48hr (IKKε, IRF3-5D). Activity is reported as relative light units (RLU), normalized to Renilla luciferase and to the stimulated treatment group. Each dot represents one independent experiment (mean of 3 technical replicates); bars show mean ± SEM. N = 3-12 independent experiments per panel. Statistics: repeated measures one-way ANOVA with Tukey’s multiple comparisons test. ns, P > 0.05; *P ≤ 0.05; **P ≤ 0.01; ***P ≤ 0.001; ****P ≤ 0.0001.

**Supplemental Figure 7: Validation of HNF4α behavior in bulk RNA-seq from HEK293T cells. (A-B)** Genome browser view of RNA-seq read coverage at the *IKBKE* (A) and *HNF4* loci in HEK293T cells (hg38). Tracks show individual replicates and are scaled identically. **(C)** Western blot staining of Vinculin, IKKε, and HNF4α in HEK293T cells expressing HNF4α, IKKε, HNF4α and IKKε, or an empty vector control. **(D)** Band intensity ratio between HNF4α and HNF4α and IKKε expressing cells in (C). Two-tailed, unpaired t-test. Each dot represents a well. *P ≤ 0.05.

**Supplemental Figure 8: Effects of overexpression of candidate secondary mediators on suppressing IFN signaling in HEK293T cells. (A-C)** ISRE promoter activity in HEK293T cells co-expressing the genes targeted by the four siRNA screen hits (*COL17A1*, *CX3CL1*, *MYO1A*, and *TRIM10*; “4 Hits”) or empty vector control, with or without the indicated stimulus. **(D-G)** ISRE promoter activity in cells expressing single siRNA screen hits or empty vector control, with or without IKKε. **(H-S)** IFNβ (H-M) and ISRE (N-S) promoter activity in cells expressing single siRNA screen hits or empty vector control, with or without IKKε. **(T-Ak)** Indicated promoter activity in cells expressing single siRNA screen hits or empty vector control, with or without indicated stimuli. Cells were co-transfected with a firefly luciferase reporter and a Renilla control and stimulated for 24hr (SenV, 5’PPP, TNF, IL-1β) or 48hr (IKKε). Activity is reported as relative light units (RLU), normalized to Renilla luciferase and to the stimulated treatment group. Each dot represents one independent experiment (mean of 3 technical replicates); bars show mean ± SEM. N = 3-12 independent experiments per panel. Statistics: repeated measures one-way ANOVA with Tukey’s multiple comparisons test. ns, P > 0.05; *P ≤ 0.05; **P ≤ 0.01; ***P ≤ 0.001; ****P ≤ 0.0001.

**Supplemental Figure 9: Infection responses in HNF4 KO animals. (A)** Schematic of chronic and acute coxsackie B virus (CVB) infection. **(B-C)** Body weights after CVB gavage after chronic (B) and acute (A) infection, shown as percent of starting weight, in WT and KO SPF mice. Symbols show the group mean ± SEM. Chronic infection n=15-16/group, IP control n=9. Acute infection n= 6-7/group, IP control n=3. Statistics: two-way repeated measures ANOVA with Šídák’s multiple comparisons test between WT, KO, and IP controls at each time point. **(D-J)** TCID50/mL of CVB in indicated tissues between HNF4 WT, KO, and IP control groups. Each dot represents one mouse. Statistics: repeated measures one-way ANOVA with Tukey’s multiple comparisons test. **(K)** Schematic of rotavirus infection. **(L)** Body weights after rotavirus gavage, shown as percent of starting weight, in WT and KO SPF mice. Symbols show the group mean ± SEM (n= 9/group). Statistics: two-way repeated measures ANOVA with Šídák’s multiple comparisons test between WT and KO mice at each time point. **(M)** Rotavirus particles per fecal pellet detected by qPCR between WT and KO mice. Each dot represents one mouse (n=9/group). Statistics: two-way repeated measures ANOVA with Šídák’s multiple comparisons test between WT and KO mice at each time point. **(N)** *S*. Tm. CFU/g of feces between WT and KO mice. Each dot represents one mouse (n=5-6/group). Statistics: two-way repeated measures ANOVA with Šídák’s multiple comparisons test between WT and KO mice at each time point. **(O-P)** *S*. Tm. CFU/g of spleen (O) or liver (P) between WT and KO mice. Each dot represents one mouse (n=5/group). Two-tailed, unpaired t-test. **(Q-V)** Histology analysis comparing intestines of WT and KO mice in indicated tissues. Each dot represents one mouse (n=7-8/group/panel). Two-tailed, unpaired t-test. ns, P > 0.05; *P ≤ 0.05; **P ≤ 0.01; ***P ≤ 0.001; ****P ≤ 0.0001.

**Supplemental Figure 10: FACs validation of magnetic bead sorting. (A-I)** Gating strategy and population information for unstained (A), single sort (B-G), and full sort (H-I) samples.

## EXPERIMENTAL MODEL AND STUDY PARTICIPANT DETAILS

### Mice

All animal procedures were approved by the Duke University Institutional Animal Care and Use Committee (protocol 103). All mice were housed and maintained at Duke University. Age and sex-matched adult mice of both sexes were used between 6-13 weeks of age across all experiments. IEC-specific HNF4α knockout mice (*Vil:Cre*^+/−^; *Hnf4a*^F/F^) and Cre-negative littermate controls (*Vil:Cre*^−/−^; *Hnf4a*^F/F^) were bred in-house under specific pathogen-free (SPF; free of *Helicobacter, Pasteurella*, and murine norovirus) conditions on a C57BL/6J background as before (27). Mice were backcrossed every 10 generations to maintain genetic homogeneity. Animals were maintained in standard bedding (Enrich-o’Cobs-The Andersons 4EB) in up to 5 animals per cage with a 12hr light/dark cycle and fed LabDiet 5053.

Germ-free (GF) C57BL/6J mice were maintained, housed, and bred in flexible-film isolators (CBC) at Duke University. Age and sex matched mice were used in all experiments. GF mice were fed ad libitum on an Teklad 2020sx chow diet and kept on a 12hr light dark cycle in up to 5 animals per cage. GF mice were housed in Alpha-Dri bedding and provided reverse osmosis water. All materials were autoclaved before being introduced to the isolators. Weekly sterility testing was performed in each isolator.

### Microbial conventionalization

GF mice were conventionalized (CV) by oral gavage with 100µL of homogenized fecal material collected from adult SPF mice. Fecal slurry was prepared with fresh, sterile phosphate buffered saline (PBS) (Gibco-14190-144) with 0.05% cysteine (Sigma-Aldrich-C1276-50G). Fecal pellets were resuspended in PBS to a 10% weight/volume slurry, manually homogenized with a plastic spreader (Fisher-14-665-230), and vortexed. Homogenization and vortexing was repeated a second time. Slurry was spun for 2min at 300g. Supernatant was collected, aliquoted, and frozen at -80°C until gavage. One batch of fecal slurry was used for all experiments. GF controls were gavaged with 100µL autoclaved drinking water.

### Cell lines

HEK293T cells (ATCC-CRL-3216) were cultured in DMEM (Thermofisher-11995065) with 10% heat inactivated fetal bovine serum (Sigma-F4135-500ML), 1X L-glutamine (ThermoFisher-10378016), and 1X Penicillin streptomycin (Millipore-Sigma-P4333-100ML) at 37°C and 5% CO₂. Cells were tested for mycoplasma quarterly.

HeLA cells were obtained by Carolyn Coyne. Cells were cultured in EMEM (Corning-10-010-CV) supplemented with 10% FBS (Sigma-F4135-500ML) and 1X Penicillin streptomycin (Millipore-Sigma-P4333-100ML) at 37°C and 5% CO₂.

### Bacterial and viral strains

*Salmonella enterica* serovar Typhimurium (strain SL1344) was gifted by Neeraj Surana. Coxsackievirus B3 and rotavirus ECwt were obtained from Carolyn Coyne and Sarah Blutt, respectively. Sendai virus-Cantell strain was obtained from ATCC (VR-907).

## METHOD DETAILS

### Infection models

#### Salmonella infection

*Salmonella enterica* serovar Typhimurium (strain SL1344) was grown overnight at 37°C in LB broth (Fisher Scientific BP1426-500) with 200µg/mL streptomycin (Millipore-Sigma-S6501-25G). After growth, bacteria was spun for 5 min at 10,000g, supernatant removed, and resuspended in sterile PBS (Gibco 14190-144). Inoculum was quantified by plating serial dilutions on LB agar (Fisher-BP1425-500) with 200µg/mL of streptomycin. Plates were kept at 37°C for 24 hours and colonies counted to calculate bacterial titre.

SPF HNF4 KO and WT age-and sex-matched littermate controls received 20mg/mouse of 0.22µm-filtered streptomycin (in sterile PBS; Millipore-Sigma-S6501-25G) by oral gavage on day −1 (100µL/mouse). On day 0, mice were gavaged with food grade 3% sodium bicarbonate in sterile PBS (Arm & Hammer Pure Baking Soda-Food lion). 30 minutes post sodium bicarbonate gavage, animals were gavaged with *S.* Tm. (3×10^8^-2×10^9^ CFU/animal) or PBS (uninfected controls). Body weight and fecal pellets were collected daily. Mice were euthanized with isoflurane (Covetrus-11695-6777-2) followed by cervical dislocation at 96 h post-infection (day 4). Liver, spleen, small intestine, and colon were collected.

### Quantification of *Salmonella* CFU

Daily fecal pellets were collected into pre-weighed 1.7mL tubes with sterile PBS. After collection, tubes were re-weighed to determine fecal and organ weights. Fecal pellets were homogenized manually with P1000 tips and vortexing. Spleen and liver samples were collected into pre-weighed 2mL tubes with ceramic homogenizing beads (Caymen Chemical-10011152). After collection samples were re-weighed and were homogenized via bead beating on a Precelly’s 24 tissue homogenizer (Bertin Technologies) for 15 seconds, 2 times at 5500rpm. Spleen and liver samples were spun for 10 min at 3000g. Fecal, spleen, and liver homogenate supernatant were serially diluted (1×10^3^-1×10^7^ for fecal pellets, pure sample-1×10^3^ for spleen and liver) and plated on LB agar with 200mg/mL streptomycin. Plates were incubated at 37°C for 24hr and counted to determine CFU. Colony counts were normalized to tissue or fecal mass to calculate CFU/g.

### Coxsackievirus B infection

SPF HNF4 KO and WT age- and sex-matched littermate controls (8-10 weeks) were orally gavaged or injected intraperitoneally (IP) with 50µL of coxsackievirus B3 (1×10^8^ PFU in PBS). Body weights were collected daily. 4 (acute infection) or 14 (chronic infection) days post inoculation, animals were euthanized with isoflurane (Covetrus-11695-6777-2) followed by cervical dislocation. Whole heart, liver, and pancreas and 1cm of each intestinal segment were collected into 1.7mL tubes, frozen in liquid nitrogen, and stored at -80°C until TCID₅₀ assays.

### Quantification of CVB by TCID_₅₀_

Heart, liver, pancreas, and intestinal segments were weighed and placed in 2mL tubes with ceramic homogenizing beads (Caymen Chemical-10011152). 29µL/mg of ice cold EMEM was added to each tube. Samples were homogenized via bead beating for 20 seconds, three times at 5500rpm on a Precelly’s 24 tissue homogenizer (Bertin Technologies). Samples were spun at 16,000g for 5 min at 4C. Supernatant was collected and placed into 1.7mL tubes and frozen untial assays.

For TCID₅₀ assays, HeLA cells were seeded at 10,000cells/well in complete EMEM (EMEM, 10% FBS, 1X Pen/strep) into 96 well plates (Cell treat-229196) and incubated at 37°C and 5% CO₂. 48hr after seeding, seeding media was removed and replaced with 225µL complete EMEM. 25µL of specified organ homogenate supernatant was added to row A. Samples were serially diluted (1:10) down the plate. Row G was left as a media control. Samples were plated in triplicate. Plates were incubated at 37°C and 5% CO₂. 48hr after dilutions, media was removed and 75µL of 1% crystal violet (Millipore Sigma-C0775-25G) was added to each well and incubated for 30min at room temperature, rotating. Crystal violet was removed and plates were gently rinsed by submerging in H_2_0, twice. TCID₅₀/mL was calculated using previously published methods (129, 130).

### Rotavirus infection

SPF HNF4 KO and WT age- and sex-matched littermate controls (8-12 weeks) were orally gavaged with 1×10^4^ ID50 of rotavirus (EcWT) from gut homogenate as previously described (131, 132). Body weights and fecal pellets were collected daily. Fecal pellets were snap frozen in liquid nitrogen until RNA extraction.

### Quantification of rotavirus genome copies

Viral RNA was extracted using the Zymo quick RNA kit (R1040) with a modified protocol. Fecal pellets were collected into 2mL tubes with ceramic homogenizing beads (Caymen Chemical-10011152). 400µL of viral RNA buffer was added to each sample tube and samples were homogenized for 1min (3 rounds, 20sec each) at 5500rpm on a Precelly’s 24 tissue homogenizer (Bertin Technologies). Samples were spun at max speed 16,000g for 2 min. 400µL of each sample was added to kit 96 well plates as per manufacturer protocol. Samples were spun for 5 min at 4000g and 500µL of kit buffer was added to each well. Plates were spun for 5 min at 16,000g, and flow through discarded. This wash step was repeated, and the well plates transferred to the kit elution plate. 50µL RNAse free H_2_0 was added to each well, incubated 2 min at room temperature, and spun for 2 min at 4000g. RNA was quantified with qubit high sensitivity RNA kit (ThermoFisher-Q32855). Rotavirus genome copies per fecal pellet were calculated by qPCR as described in the RT-qPCR section.

### Histology

Small intestine and colon samples from HNF4 KO and WT mice infected with *S*. Tm. were collected into ice cold PBS (Gibco 14190-144) with 10μM Y-27632 dihydrochloride (ApexBio A3008-50). Samples were flushed with Carnoy’s solution (60% EtOH [VWR V1001], 30% chloroform [Sigma-Aldrich 1024441000], and 10% glacial acetic acid [Fisher Scientific A38-500]), flayed open, and swiss rolled. Once rolled, samples were fixed and stored in 10% formalin (Fisher Scientific-22-050-259). Samples were paraffin-embedded, sectioned, and stained with hematoxylin and eosin by the Duke Pathology Biorepository and Precision Pathology Center Rresearch Histology Laboratory. Slides were examined by a board-certified veterinary pathologist blinded to experimental group. For the small intestine, villus length and crypt depth were measured (µm) in each available section (duodenum, jejunum, ileum). Measurements were obtained only in areas with exceptional orientation in which the entirety of a crypt and villus could be identified. Villus length was divided by crypt depth to calculate villi/crypt ratios. Villus blunting/fusion and inflammation were scored holistically along the entire small intestine ordinally between 0-4 (correlating to none, minimal, mild, moderate, or severe). Villus blunting/fusion and inflammation scores were summed for a severity score. Colonic epithelial damage and inflammation were similarly scored and summed for a colon severity score and crypt depth was similarly measured in both the proximal colon and distal colon. Villus length and crypt depth measurements were taken at 10 semi-randomly selected locations and averaged. Locations were inclusive of proportional representation of lesion regions.

### Isolation of IECs from jejunum

IECs were isolated as previously described with modification (133). Small intestines were removed and the middle half of the intestine was collected and flushed with ice cold epithelial buffer (EB) (5.6mM disodium phosphate, 8mM monopotassium phosphate, 96.3mM sodium chloride, 1.6mM potassium chloride, 43.4mM sucrose, and 54.9mM d-Sorbitol) with 10μM Y-27632 dihydrochloride. Samples were flayed longitudinally and washed in fresh epithelial buffer by shaking. Intestinal sections were cut into ∼1cm pieces and incubated in IEC extraction buffer (epithelial buffer with 3mM EDTA) for 15min at 4°C, rocking. Segments were transferred to EB and shaken for 2 min at 150 beats per min (bpm). Tissue was transferred to new IEC extraction buffer and incubated for 45min at 4°C, rocking. Fragments were moved to fresh EB and shaken for 4 min at 150bpm. Remaining tissue pieces were removed and isolated crypts and villi were spun at 300g for 5 min at 4°C and supernatant removed. Pellets were processed for scRNA- or BULK RNA-seq separation as below

### Dissociation of jejunal IECs for scRNA-seq

Isolated jejunal pellets (described in the isolation of IECs from jejunum section) were resuspended in dissociation buffer (EB, 10mg/mL cold protease [Sigma P5380-250MG], 2.5mg/mL DNASE [Sigma DN25-100MG]). Samples were rocked at 4°C and manually shaken (vigorously) for 10 seconds every 2-3 min. Dissociation was monitored until cells were ∼90% dissociated, never exceeding 27min. After dissociation was complete, 5mL of FACS buffer (DMEM [Sigma-Aldrich- D6429-6X1L], 10% FBS, 10μM Y-27632 dihydrochloride) was added and samples were spun at 300g for 5 min at 4°C. Supernatant was discarded and samples were resuspended in fresh FACS buffer and filtered into a 5mL falcon tube with a filter cap (Corning-352235). Cells were counted in 1:1 trypan blue using a TC20 automated cell counter (BioRad). Only cells with a diameter of 8-11µm were counted. Cells were spun at 300g for 5 min at 4°C and supernatant discarded. Samples were resuspended in 1mL FACS and transferred to 1.7mL tubes. Single color and fluorescent minus one (FMO) controls were pooled. Samples and controls were pelleted at 300g for 2 min at 4°C. Supernatant was decanted to ∼100µL and cells were stained with -EpCAM-eFluor 450 (5µL/1×10^6^ cells; eBiosciences, 48-5791-82), PE/Cyanine 7 anti-mouse TER-119/Erythroid cells (0.3µL/1×10^6^ cells; BioLegend-116221), PE/Cyainine 7 anti-mouse CD45 (0.3µL/1×10^6^ cells;BioLegend, 103114), and PE/Cyanine 7 anti-mouse CD31 (0.3µL/1×10^6^ cells;BioLegend; 102418) for 30 min at 4°C, rocking. Cells were washed 3x in 1mL FACS and spun at 300g for 2 min each at 4°C. Following the last wash, cells were resuspended in 500µL FACS. 5nM Helix NP Green (BioLegend-425303) was added to each sample and incubated for 5 min on ice. Samples were filtered over a 70µm strainer (Celltreat-229484) and FACS sorted.

### FACs sorting and scRNA-seq library preparation

After exclusion of debris and doublets, live, dump-negative, EpCAM⁺ cells were sorted on a SH800Z cell sorter (Sony) into FACS buffer. Gates were set with unstained, single-stain, and EpCAM FMO controls (Figure S1A-G). Libraries were prepared for scRNA-seq at the UNC Advanced Analytics Core using the 10x Genomics’s 3’ v3.1 scRNA-Seq workflow with CellPlex cell multiplexing oligo (CMO). The scRNA-Seq workflows were performed at the AAC according to 10x’s specifications.

### scRNA processing and analysis with Seurat

For 3 GF and 3 CV male biological replicates 10X 3’ v3.1 Gene Expression (GEX) and Cell Multiplexing (CMO) sequencing reads were generated with a confidence threshold of 0.7 and expectation of 10,000 cells. Sequencing reads were mapped using Cell Ranger (10X Genomics) to mm10-2020-A genome. CV and GF conditions were merged separately using Seurat (v5.0.0+) pipelines from the Satija lab (https://satijalab.org/seurat/articles/). Cells were subset to include cells with > 200 and < 6,000 features and < 25% mitochondrial transcripts. For each condition, data was scaled separately to regress percent of mitochondrial reads. To integrate the two conditions, standard pairwise integration pipelines were followed using 2000 features and 1:20 dims for integration anchors (https://satijalab.org/seurat/articles/integration_mapping.html) and, PCA reduction and a resolution of .6 for cluster segregation. Cell proportion significance was tested between conditions using propeller from the Speckle package.

FindAllMarkers command was used to generate cluster-specific markers with < .05 padj and >.25 log2FC. AggregateExpression was used to pseudobulk counts per replicate and cluster. Enterocyte subset was characterized as cluster 0-6. DESeq2 was run to contrast the CV versus GF condition exporting cluster specific raw log2FC. When required <.05 p-adjusted and an absolute value log2FC >.5 between conditions was used to call significant genes. A matrix of binary cluster specific DEGs between CV/GF was hierarchically clustered using Cluster 3.0 correlation uncentered with average linkage. Gene set enrichment analysis (GSEA) was run from the clusterprofiler package using Hallmark gene sets from msigdb and Reactome gene sets using ReactomePA. For gene lists, Metascape (Metascape.org) was used with default parameters and mouse as the analysis species.

Gene lists for genes coincidently positive (Up) or negative (Down) for FindAllMarkers and DEGs from the CV/GF enterocyte contrast were used for motif enrichment using cisTarget mm10 refseq-r80 10kb_up_and_down_tss.mc9nr.feather and mm10 refseq-r80 500bp_up_and_100bp_down_tss.mc9nr.feather motif positions

### Virus search in single cell sequencing datasets

cDNA reads from the above 10X scRNAseq data were mapped to the Mus musculus genome assembly GRCm38 using STAR (134) version=2.7.10b. Unmapped reads were assembled de novo using SPAdes (135) version: 4.0.0. Assembled transcripts were aligned to protein sequences from the NCBI clustered nr database using DIAMOND (136) version 2.0.15.153 with the --very-sensitive setting and taxonomically annotated using the resulting target sequence with the highest bit score. Sequences prospectively annotated as viruses based on protein sequence similarity were subsequently aligned to all nucleotide sequences from the NCBI nt database using BLAST (137) version 2.12.0+ and updated if the bit score for the best nucleotide alignment exceeded that of the protein alignment.

### Quantification of transposable elements in single cell sequencing datasets

cDNA reads from the above 10X scRNAseq data were mapped to the *Mus musculus* genome assembly GRCm39 and UCSC mm39 RepeatMasker annotation gtf using STAR (134) version=2.7.10b with options --outFilterMultimapNmax 100 --winAnchorMultimapNmax 200. Transposable elements were quantified at the locus level using Telescope (138) version 1.0.3 and differential expression analysis was performed with DESeq2 (139) version 1.52.0. Enrichment of TE families among differentially expressed loci was quantified by one-sided hypergeometric enrichment of each TE family among the significant loci compared to all tested loci with Benjamini-Hochberg correction.

### Virus PCR assays

Fecal pellets collected from the GF and SPF mouse colonies used in this study were screened by PCR (IDEXX BioAnalytics) for the following viruses: Ectromelia, EDIM, MHV, MNV, MVM, MPV, TMEV, Hantaan, LCMV, LDEV, MAV1, MAV2, MCMV, MTV, Polyoma, PVM, REO3, Sendai, and K virus. All assays were negative.

### Bulk RNA-seq of HEK293T cells

HEK293T cells were seeded at 50,000 cells/well in 24 well plates (cell treat-229124) in complete EMEM. 24hr post seeding HEK293T cells were transfected with an empty vector, HNF4α, IKKε, or HNF4α + IKKε using 25ng of the respective plasmids, Opti-MEM reduced serum media (ThermoFisher-31985062), and Fugene 6 transfection reagents (Promega-E2691) according to manufacturer’s protocols. RNA extraction was performed using the ISOLATE II RNA Mini Kit (Bioline-BIO-52073) according to the manufacturer’s protocol. RNA was quantified by qubit broad range RNA kit (ThermoFisher-Q10210) according to the manufacturer’s protocol. Library preparation and sequencing were performed by the Duke Sequencing and Genomic Technologies. RNA was quantified using (Qubit™ RNA HS Assay Kit, cat #Q32852) on (Varioskan Lux). From each sample, (100) ng of input RNA was used for library preparation with (KAPA mRNA HyperPrep Kit, cat #KK8581) following manufacturer instructions with (16) cycles of PCR. Final libraries were quantified with (Qubit DNA HS, cat #Q32854) on (Varioskan Lux) and equimolar libraries were pooled. The final pool was quantified using (Qubit DNA HS, cat #Q32854) on (Varioskan Lux), diluted to (130pM+1%phiX), and loaded onto (Novaseq X plus) using (10B 300, cat #20085594) to generate (151PE) reads for a target read depth 110M reads/sample.

### Computational Processing HEK293T RNA-seq

HEK293T sequencing reads were mapped to GRch38 using STAR (2.7.10b) and counts were generated with featureCounts using annotation Homo_sapiens.GRCh38.110.chr.gtf. Differential gene calls were generated using Wald contrasts between Ikke/Ctrl, Hnf4a/Ctrl, Hnf4aIkke/Hnf4a and Hnf4aIkke/Ikke and interaction analysis using likelihood ratio test (LRT) between Ikke status and Hnf4a status. Clustering was performed using Cluster 3.0 correlation uncentered and average linkage on row z-scored normalized counts for genes differential in at least one contrast. For gene lists, Metascape (Metascape.org) was used with default parameters and human as the analysis species.

### Bulk RNA-seq single cell dissociation and magnetic bead isolation of mouse IECs

HNF4 KO and WT mice small intestines were collected and measured. Intestinal length was divided by 30 and corresponding jejunal sections 10-17 were collected, as described previously (127). IECs were isolated as above. Pellets were resuspended in 10mL TrypLE express enzyme (ThermoFisher-12604013) supplemented with 2.5mg/mL DNASE (Sigma DN25-100MG) and 10μM Y-27632 dihydrochloride. Samples were shaken at 30sec at 150bpm every 3 minutes. Dissociation was monitored until ∼90% of cells were in single cell suspension, never extending past ∼27min. Cells were filtered over a 70µM filter into 40mL FACS and inverted 10x. Samples were then filtered over a 40µm filter (Cell Treat-229481) and spun for 10min at 600g at 4°C. Supernatant was decanted and cells were resuspended in 5mL FACS. Samples were counted in 1:1 trypan blue using a TC20 Automated cell counter (BioRad). Only cells with a diameter of 8-16µm were counted. Samples were spun for 5 min at 600g at 4°C, decanted, resuspended in 1mL EB and transferred to a 1.7mL tube. Cells were washed 2x in 1mL EB and decanted. Cells were washed a final time and supernatant was aspirated completely. Cells were magnetically labeled and dead cells removed using the dead cell removal kit (Miltenyi Biotec-130-090-101) according to the manufacturers protocol using a quadroMACs separator (Miltenyi Biotec-130-042-108). Following live cell enrichment, cells were sequentially labeled for CD45 (Miltenyi Biotec-130-052-301) and EpCAM (Miltenyi Biotec-130-105-958) and the CD45^-^, EpCAM^+^ populations were collected. Cells were pelleted at 300g for 10min at 4°C and supernatant aspirated completely. Validation of population enrichment was performed by FACS on a FACSCanto machine (BD). Pellets were then subjected to RNA extraction using the ISOLATE II RNA Mini Kit (Bioline-BIO-52073) according to the manufacturer’s protocol. RNA was quantified by qubit broad range RNA kit (ThermoFisher-Q10210) according to the manufacturer’s protocol.

### Bulk RNA-seq of mouse IECs

Library preparation and sequencing were performed by the Duke Sequencing and Genomic Technologies. RNA was quantified using (Quant-it BR RNA, cat #Q10213) on (Varioskan Lux). From each sample, (200) ng of input RNA was used for library preparation with (Watchmaker mRNA prep, cat #7BK0001-096) following manufacturer instructions with (15) cycles of PCR. Final libraries were quantified with (Qubit DNA HS, cat #Q32854) on (Varioskan Lux) and equimolar libraries were pooled. The final pool was quantified using (Qubit DNA HS, cat #Q32854) on (Varioskan Lux), diluted to (130pM+1%phiX), and loaded onto (Novaseq X plus) using (25B 300, cat #20104706) to generate (151PE) reads for a target read depth 50M reads/sample.

### Processing of Mouse IEC RNA-seq

Mouse IEC Salmonella infection experimental replicate reads were mapped to GRCm38 using STAR (2.7.11b) and counts were generated with featureCounts using annotation Mus_musculus.GRCm38.80.gtf. Differential gene calls were generated using Wald test contrasts between UIKO/UIWT, IKO/IWT, IWT/UIWT, IKO/UIKO and interaction analysis using likelihood ratio test (LRT) and Wald interaction test between Hnf4a status and Infection status. Clustering was performed using Cluster 3.0 correlation uncentered and average linkage on row z-scored noramlized counts for genes differential in at least one contrast. Gene set enrichment analysis (GSEA) was run from the clusterprofiler package using Hallmark gene sets from msigdb and Reactome gene sets using ReactomePA. For gene lists, Metascape (Metascape.org) was used with default parameters and mouse as the analysis species.

HEK293T sequencing reads were mapped to GRch38 using STAR (2.7.10b) and counts were generated with featureCounts using annotation Homo_sapiens.GRCh38.110.chr.gtf (Ensembl.org). Differential gene calls were generated with DESeq2 using Wald test contrasts between Ikke/Ctrl, Hnf4a/Ctrl, Hnf4aIkke/Hnf4a and Hnf4aIkke/Ikke and interaction analysis using likelihood ratio test (LRT) between Ikke status and Hnf4a Status. Clustering was performed using Cluster 3.0 correlation uncentered and average linkage on row z-scored noramlized counts for genes differential in at least one contrast. For gene lists, Metascape (Metascape.org) was used with default parameters and human as the analysis species.

### Plasmids and cloning

The IFNβ, NF-κB, and ISRE luciferase reporters, renilla luciferase reporter, IKKε, and IRF3-5D plasmids were gifts from Stacy Horner. The TREH promoter luciferase reporter was a gift from Jesper Troelsen (68).

The HNF4α^AAA^ plasmid was created using the HNF4α plasmid and NEB’s site directed mutagenesis protocol and the KLD enzyme reaction mix (NEB #M0554). Primers used for site-directed mutagenesis:

F-CAGCAAAGAGGAACCAGTGCCG,
R-CTGCCACCACGCACTGCCG.

Plasmid mutation was validated by sequencing through Plasmidaurus. Predicted structures of WT HNF4α and mutant HNF4α^AAA^ were generated with AlphaFold and compared in ChimeraX (version 1.12) by RMSD of defined regions.

Human cDNA constructs for *COL17A1* (HsCD00860488), *CX3CL1* (HsCD00039574) were obtained from DNAsu. PcDNA3, *COL17A1*, *CX3CL1* plasmid fragments were prepared for Gibson cloning with the primers listed below. Fragments were joined via Gibson cloning following the manufacturer’s protocol (NEB-E5510S) to generate *COL17A1* and *CX3CL1* plasmids in the pcDNA3 backbone driven by the CMV promoter. Gibson reactions were then transformed into GC5 competent cells (Genesse-42-652) following the NEB transformation protocol. Transformed bacteria were plated on LB agar (Fisher scientific-BP1425-500) supplemented with 100µg Carbenicillin sodium salt (Fisher Scientific-10177012). Plasmids were extracted by ZymoPURE mini- or maxi-prep kits following the manufacturer’s protocol (Thomas Scientific-1159V20, VWR-77001-470). Plasmids were validated by sequencing through Plasmidaurus. Primer sequences used:

COL17-F-cggccgccactgtgctggattcaCAACGGCTTGACAGCAATAC
COL17-R-tgtggtggaattctgcagatgccgccaccATGGATGTAACCAAGAAAAACAAAC CX3CL1-F-cggccgccactgtgctggattcaCACGGGCACCAGGACATATG
CX3CL1-R-tgtggtggaattctgcagatgccgccaccATGGCTCCGATATCTCTGTCG pcDNA3-F-TGGTGGCAAGCTTAAGTTTAAAC
pcDNA3-R-TAGGGATCCGAATTCTGC

Custom plasmids containing *MYO1A* and *TRIM10* driven by the CMV promoter were purchased from VectorBuilder. Plasmids were extracted by ZymoPURE mini- or maxi-prep kits following the manufacturer’s protocol (Thomas Scientific-1159V20, VWR-77001-470). Plasmids were validated by sequencing through Plasmidaurus.

### HOMER Motif Analysis

Transcription factor motif enrichment was performed using HOMER (Hypergeometric Optimization of Motif EnRichment; v3.12]) (140) Putative HNF4α binding sites in whole IFNβ and NF-κB reporter plasmid sequences were searched with against allHomer.motifs.

### Luciferase reporter assays

HEK293T cells were seeded at 10,000 cells/well in 96 well plates (Celltreat-229196) in complete DMEM. 24hr post plating, cells were transfected with 50ng firefly luciferase reporter, 1ng Renilla luciferase control, and 5ng of designated overexpression plasmids using Fugene6 (total ng plasmids x 3/1000) (Promega- 229196) in Opti-MEM reduced serum media (ThermoFisher-31985070).

24hr after transfection, unless transfected with IKKε or IRF3-5D, cells were stimulated with human recombinant TNFα (5ng/mL; ThermoFisher-4408), human recombinant IL-1β (5ng/mL, SinoBiological-10139-HNAE), human IFNλ1 (20ng/mL, Peprotechl-300-02L), or human IFNβ (10ng/mL, ThermoFisher-300-02BC-20UG) in complete DMEM. Where indicated, cells were pre-treated for 1hr with ruxolitinib (8µM; MedChem Express-HY-50856) before IFN treatments were added. Ruxolitnib was present in the media for the remainder of the experiment.

For Sendai Virus treatment (Cantell stain, ATCC-VR-907), cells were given 200µL of complete media with 1:20 dilution of SenV. Before being added to media, SenV was incubated in 20µL Opti-MEM reduced serum media for 15 min at room temperature.

5′-triphosphate RNA was prepared in house and transfected into indicated cells (1µg/µL stock) using the Mirus Bio TransIT-mRNA Transfection Kit (VWR-10767-106) following the manufacturer’s protocol.

Cells were assayed 24hr after stimulation (TNFα, IL-1β, SenV, 5′PPP, IFNs) or 48hr after transfection (IKKε, IRF3-5D). Firefly and Renilla luminescence were measured with a Dual-Glo luciferase assay system (Promega-E2920) and read on a Synergy 2 microplate reader (BioTek) according to the manufacturer’s protocol. Normalization of the signal was carried out according to the manufacturer’s protocol.

### Generation of 5’-Triphosphate RNA

NeoPoluUA and NeoPoluUS primers (1µg/µL stocks) were annealed with the following conditions: 95°C for 10min, 85°C for 1min, 75°C for 1min, 65°C for 1min, 55°C for 1min, 45°C for 1min, 35°C for 1min, 25°C for 1 min, followed by a 4°C hold. Once annealed, *in vitro* transcription was performed using the Ambion MegaShortScript T7 Kit (ThermoFisher-AM1354) with 1µL of annealed primers, 2µL of the kit reaction buffer, ATP, CTP, GTP, UTP, and enzyme solutions, and 7µL nuclease free water. The solution was incubated at 37°C for 3hrs followed by 1µL treatment of TURBO DNase (ThermoFisher-AM1907) for 15 min at 37°C. After DNase treatment, 115µL of nuclease free H_2_0 and 15µL of Ammonium Acetate Stop solution (MegaShortScript T7 kit) was added and mixed thoroughly. 2 volumes of 100% EtOH was added and chilled at -20°C for 15min. Samples were centrifuged at max speed for 15min at 4°C. Supernatant was discarded and resuspended in nuclease free H20. RNA was quantified by qubit broad range RNA kit (ThermoFisher-Q10210) according to the manufacturer’s protocol. Samples were aliquoted at 1µg/µL and stored at -80°C until transfection. Oligo sequences used:

NeoPolyUS-TAATACGACTCACTATAGGCCATCCTGTTTTTTTCCCTTTTTTTTTTTCTTTTTTTTTTTTTTTT TTTTTTTTTTTTTTTTTTCTCCTTTTTTTTTCCTCTTTTTTTCCTTTTCTTTCCTTT
NeoPolyUA-AAAGGAAAGAAAAGGAAAAAAAGAGGAAAAAAAAAGGAGAAAAAAAAAAAAAAAAAAAAAA AAAAAAAAAAAAGAAAAAAAAAAAGGGAAAAAAACAGGATGGCCTATAGTGAGTCGTATTA

### siRNA screen

siRNAs targeting the 28 genes upregulated by HNF4 and relevant controls were ordered as OnTargetPLUS custom smart pooled siRNA from Horizon. HEK293T cells were plated at 10,000cells/well in 96well plates as before. 24hrs post seeding, cells were transfected with one pooled siRNA target per well (100nM), in triplicate using RNAi Max (0.5µL/well; ThermoFisher-13778075) in Opti-MEM reduced serum media (10µL/well; ThermoFisher-31985070). 24hr post siRNA transfection, cells were co-transfected with the IFNβ reporter, Renilla control, HNF4α, and IKKε, and luciferase activity was measured as above. The entire screen was conducted 5 times independently (3 technical replicates per repeat), and results reported cumulatively.

### Quantitative RT-PCR

Normalized RNA from viral extraction, organoids, or HEK293T cells was reverse transcribed to cDNA using the iScript cDNA synthesis kit according to the manufacturer’s protocol (BioRad-1708891). RNAse free H_2_0 was used to dilute cDNA 1:10. TB Green® Premix Ex Taq™ II FAST qPCR master mix was used to quantify transcript levels on QuantSudio 6 Pro-real time system (ThermoFisher A43180). Expression of *ISG15*, *IFNL1*, and *IFNB1* (HEK293T), or *Isg15* (organoids), was normalized to *GAPDH*/*Gapdh* by the ΔΔCt method and expressed relative to control groups. Rotavirus genome copies per fecal pellet were quantified via g block standard curve as previously described (141–143). Oligo sequences used:

Rotavirus

F: CTTCTGGACTACGACAGACTATTT,
R: GATGTTTGATCGGTTCGTTGTG

ISG15 (Human)

F-GCGAACTCATCTTTGCCAGTA
R-CCAGCATCTTCACCGTCAG

IFNL1 (Human)

F-CTTCCAAGCCCACCACAACT
R-GGCCTCCAGGACCTTCAGC

IFNB1 (Human)

F-GCCAGGAGGTTCTCAACAAT
R-CTTTGCTATTTTCACACAAGATTCA

GAPDH (Human)

F-AAGGTGAAGGTCGGAGTCAAC
R-GGGGTCATTGATGGCAACAATA

Isg15 (mouse)

F-TGCCTGCAGTTCTGTACCAC
R-AGTGCTCCAGGACGGTCTTA

Gapdh (Mouse)

F-GGAGAAACCTGCCAAGTATG
R-GGAGTTGCTGTTGAAGTCG

Rotavirus gblock sequence ATTTAATGCTTTTCAGTGGTCGTTGCTCAAGATGGAGTCAACTCAGCAGATGGCTAG TTCTATTATAAATTCTTCATTCGAGGCTGCTGTAGTTGCAGCGACATCAACGCTGGAGCTGAT GGGCATTGAATATGATTACAATGAGGTGTTCACCAGAGTCAAGAGTAAATTTGAATTCGTCAT GGACGACTCCGGCGTTAGGAACAACTTGATGGGCAAAGCGTTTACCATAGATCAGGCTCTA AACAACAGATTCAACTCAGCTATGAGGAACAGAAATTGGATGTGTGACTCAAAAACTATTTCC AAGCTGGATGAAGATGTGAACAAATTGAGGATGATGTTGTCTTCCAAAGGTATAGATCAAAA AATGAGAGTCCTGAATTCATGCTTCAGTGTAAAGAGGATACCAGGTAAATCATCTTCAATAATT AGATGCACAAGACTGATGCGTGATAAACTAGAGAGAGGAGAAGTAGAAGTGGACGACAGCT TTGTCGAGGAGAGGATGGAAACTGACACTATAGACTGGAAATCACGCTACGAGCAATTGGA GAAAAGGTTTGAGACATCGAAGCAAAAAACCACAGAGAAATACAACGCATGGGCTGCAAAA GCTAAGAAAGCTAATGAAAACCTAAACGCTCTGCAATCGGTCATCACTCAACAACAAAAAAC TATAATGGAGCTACAAGAGCAGAACAGACGTATCGAGGCTGAATTACAGGCGAAGCTGTCAT CGGTGACCGGATCTATTGAATGGTTCCTGA

### Intestinal organoids

Organoid 2x basal media was prepared fresh weekly. 2X basal media was comprised of Advanced DMEM F12 (ThermoFisher-12634010) containing 2X Glutamax (ThermoFisher-35050061), 2X B27 -Vitamin A (ThermoFisher-12587010), 2X N2 (ThermoFisher-17502048), 20mM HEPES (ThermoFisher-H0887-100ML), 2mM N-Acetyl-L-cysteine (MBio-219460305), 200µg/mL primocin (InvivoGen-ant-pm-1), and 1x Pen/Strep (ThermoFisher-P4333-100ML).

Complete organoid media was prepared fresh daily. Complete organoid media contained 50% organoid 2x basal media, 50% Advanced DMEM F12, 250µg/mL recombinant murine Wnt3a (Peprotech-315-20), 100µg/mL recombinant murine Noggin (Peprotech-250-38), 100µg/mL recombinant murine R-Spondin1 (Peprotech-315-32), and 50µg/mL recombinant murine EGF (Peprotech-315-09).

Jejunal crypts from intestinal segments 10-17 were isolated from HNF4 KO and WT littermates (6-13 weeks) as above. Intestinal segments were flushed with ice cold EB with 10μM Y-27632 dihydrochloride. Samples were flayed longitudinally and washed in fresh epithelial buffer by shaking 10x, gently. Intestinal sections were cut into ∼1cm pieces and incubated in IEC extraction buffer (epithelial buffer with 3mM EDTA) for 15min at 4°C, rocking. Segments were transferred to EB and shaken for 2 min at 150bpm, gently. Tissue was transferred to new IEC extraction buffer and incubated for 45min at 4°C, rocking. Fragments were moved to fresh EB and shaken for 4 min at 150bpm, vigorously. Remaining tissue pieces were removed and crypts were counted. 5 crypts/µL were embedded in 100µL matrigel mixture/well (60% matrigel [Corning-356255]l, 40% complete organoid media) in 24 well plates (Celltreat-229124). Complete media was changed on days 2 and 4. On day 4, organoids were stimulated with PBS, high molecular weight poly I:C (100µg/mL, InvivoGen-tlrl-pic), IFNβ (10ng/mL, SinoBiological-50708-MCCH), or IFNλ [10ng/mL, SinoBiological-51305-M08H]. For poly I:C treatments, the Mirus Bio TransIT-mRNA Transfection Kit (VWR-10767-106) was used at 2µL of kit reagent and 2µL of kit boost per 1mL media. Reagent and boost were added to poly I:C for 3 min before being combined into media. Media was then added to wells. IFNβ or IFNλ were added directly to complete media. Organoids were harvested 24hr post stimulation for RNA extraction and qPCR as described above. Each well was treated as a technical unit and wells were derived from 4-7 animals per genotype.

### Western blotting

HEK293T cells were plated at 50,000 cells/well in 24 well plates and transfected with indicated plasmids at 75ng/plasmid as described above. Cells were scraped from plates and lysed by vortexing in a modified radioimmunoprecipitation assay (RIPA) buffer supplemented with 1X protease inhibitor (cell signaling-5872) and postnuclear lysates were isolated by centrifugation. Protein was quantified via BCA Protein Assay Kit according to manufacturer’s protocol (ThermoFisher-23227-thermo). Normalized protein (between 2 and 15 μg) was resolved by sodium dodecyl sulfate (SDS)/polyacrylamide gel electrophoresis (PAGE) (Biorad-4568086), transferred to nitrocellulose in a 25 mM Tris-192 mM glycine-0.01% SDS buffer using the Trans-Blot Turbo Transfer System (BioRad). Membranes were stained with Ponceau S (ThermoFisher-A40000279) and then blocked in 5% bovine serum albumin (Sigma-F4135-500ML) or 5% milk (Kroger) in Tris-buffered saline containing 0.01% Tween-20 (TBST) for 1hr at room temp. Membranes were then incubated in primary antibodies overnight at 4°C, rocking. The next day, membranes were washed in TBST 3x for 5 min at room temperature. Membranes were incubated in horseradish peroxidase (HRP)–conjugated antibodies in 5% BSA for 3hr at room temperature and washed in TBST 3x. Finally membranes were treated with Clarity Western ECL substrate (Bio-Rad-170-5061) before imaging on a ChemiDoc Imaging system (BioRad). Bands were quantified in ImageJ and HNF4α was normalized to vinculin. The following antibodies were used for immunoblotting: Recombinant anti-HNF4α (1:1000 in 5% milk; Abcam-ab181604), anti-Vinculin (1:1000 in 5% BSA; Cell Signaling Technologies-# 13901S), anti-IKKε (1:1000 in 5% BSA; Cell Signaling Technologies-3416S), and goat anti-rabbit HRP (1:10,000 in 5% BSA, Boster Biological Technology, BA-1054-0.5).

### Immunofluorescence and confocal microscopy

HEK293T cells were plated at 50,000cells/well in 24 wells on 15mm round glass coverslips (Celltreat-229172). 24hr after seeing, cells were transfected with 75ng HNF4α, HNF4α^AAA^, IKKε, or combinations as above. 48hr after transfection, media was removed and cells were fixed in 4% PFA (Sigma-158127-500G) for 20min at room temperature. Cells were washed 3x in PBS(Gibco-14190-144) and blocked in TBST with 1:200 Anti-mouse CD16/CD32FC block (Invitrogen, 14-0161-82), and 2% normal goat serum (Jackson ImmunoResearch Laboratories-005-000-121) for 1hr at room temp. Cells were incubated in primary antibodies overnight in blocking solution (1% BSA and 0.5% Triton-X 100 [Fisher Scientific-BP151-100], and 2% normal goat serum in 1X PBS) at 4°C in a humidity chamber. The next day, samples were washed 3x with TBST then stained with secondary antibodies in blocking solution for 3hr at room temperature in the dark. Cells were washed 3x with TBST then stained with a nuclear stain for 15 min in blocking solution in the dark. Cells were washed 3x in Tris-Buffered Saline (TBS; 20mM tris base [Fisher Scientific-BP152-500] and 150mM sodium chloride) and coverslips removed from wells. Coverslips were mounted on glass slides (Fisherbrand-12-550-15) with Prolong Gold Antifade Mounting Media (Invitrogen- P10144) and sealed with clear nail varnish (Amazon-C450B). Images were captured with an Andor iXon888 EMCCD camera attached to the Dragonfly 505 spinning disk unit (25um or 40um pinhole) on a Leica DMi8 inverted stand using the Leica HC PL APO 63x water objective. The sample was imaged with 405nm laser excitation and 460/50 emission filter, 488nm laser excitation and 525/50 emission filter, 561nm excitation and 600/50 emission. The motorized stage has linear encoding (Applied Scientific Instrumentation-ASI) and z-stacks were collected with piezo z-stage with a 300 micron range. Hardware was controlled with Andor Fusion software version 2.7. Identical acquisition settings were used across conditions and processed in Fiji. Scale bars are indicated on each image.

Antibodies used were: Goat anti-mouse 488 (1:1000, Invitrogen-A11001, Goat anti-rabbit 647 (1:1000, Invitrogen-A21244), Goat anti-rabbit IgG (1:1000 of 200μg/mL, JacksonImmunoResearch-111-005-003), Goat anti-mouse IgG (1:1000 of 200μg/mL, JacksonImmunoResearch-115-005-044), mouse anti-human HNF4α (1:1000, Invitrogen-MA1-199), rabbit anti-human IKKε (1:1000, Cell Signaling Technologies-3416), Hoechst 33342 (1:700, Life Technologies-H3570).

## QUANTIFICATION AND STATISTICAL ANALYSIS

Statistical analyses and graphing were performed in GraphPad Prism 11 for all non-sequencing experiments. Sample sizes (n) and the definitions of replicates and statistical tests are given in the figure legends. Significance thresholds were: ns, P > 0.05; *P ≤ 0.05; **P ≤ 0.01; ***P ≤ 0.001; ****P ≤ 0.0001.

