## Supplemental Figures for "Intestinal epithelial differentiation factor HNF4α restrains homeostatic interferon signaling and is required for its full induction during *Salmonella* infection"

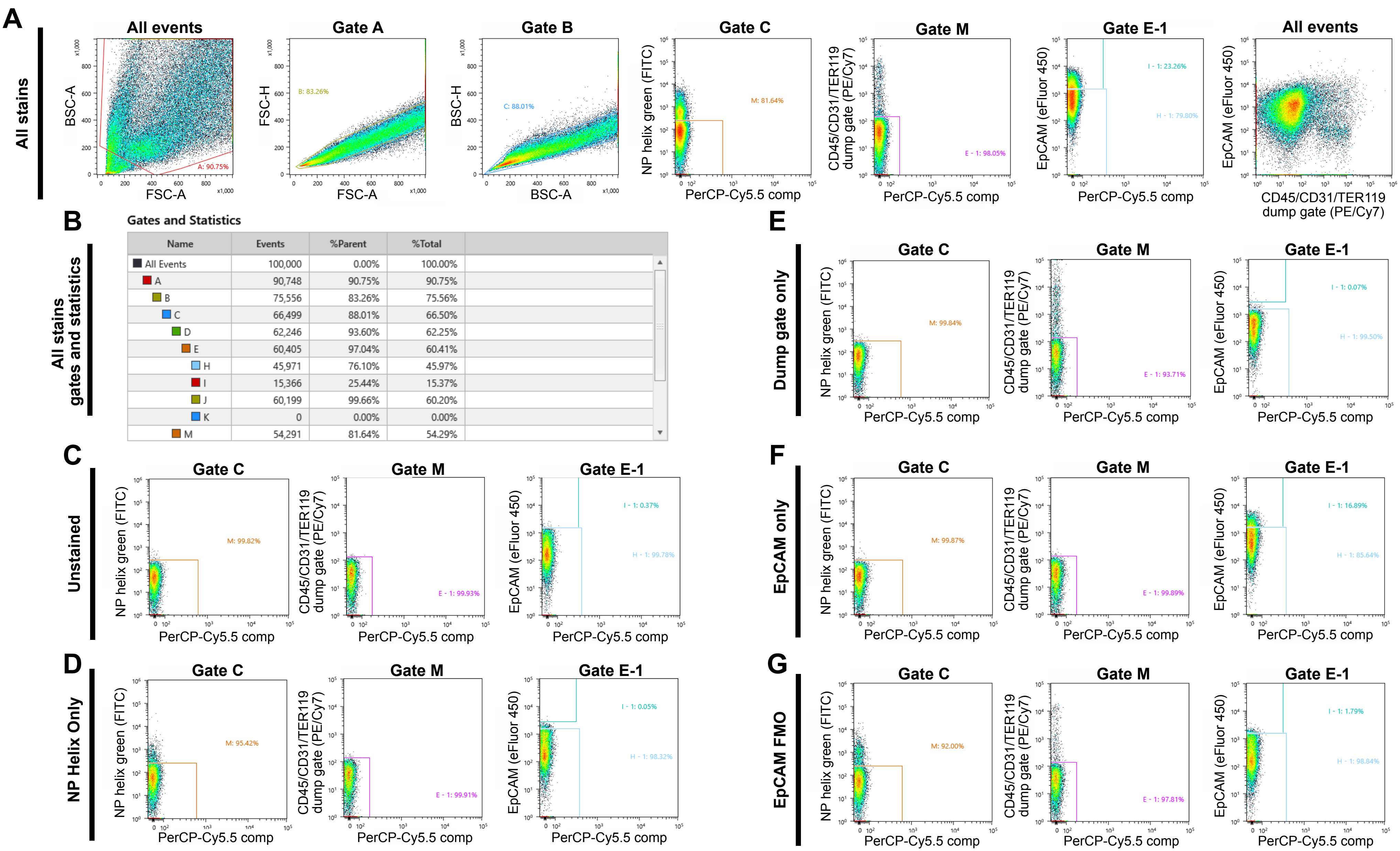

Gates and Statistics

| Name | Events | %Parent | %Total |
|---|---|---|---|
| All Events | 100,000 | 0.00% | 100.00% |
| A | 90,748 | 90.75% | 90.75% |
| B | 75,556 | 83.26% | 75.56% |
| C | 66,499 | 88.01% | 66.50% |
| D | 62,246 | 93.60% | 62.25% |
| E | 60,405 | 97.04% | 60.41% |
| H | 45,971 | 76.10% | 45.97% |
| I | 15,366 | 25.44% | 15.37% |
| J | 60,199 | 99.66% | 60.20% |
| K | 0 | 0.00% | 0.00% |
| M | 54,291 | 81.64% | 54.29% |

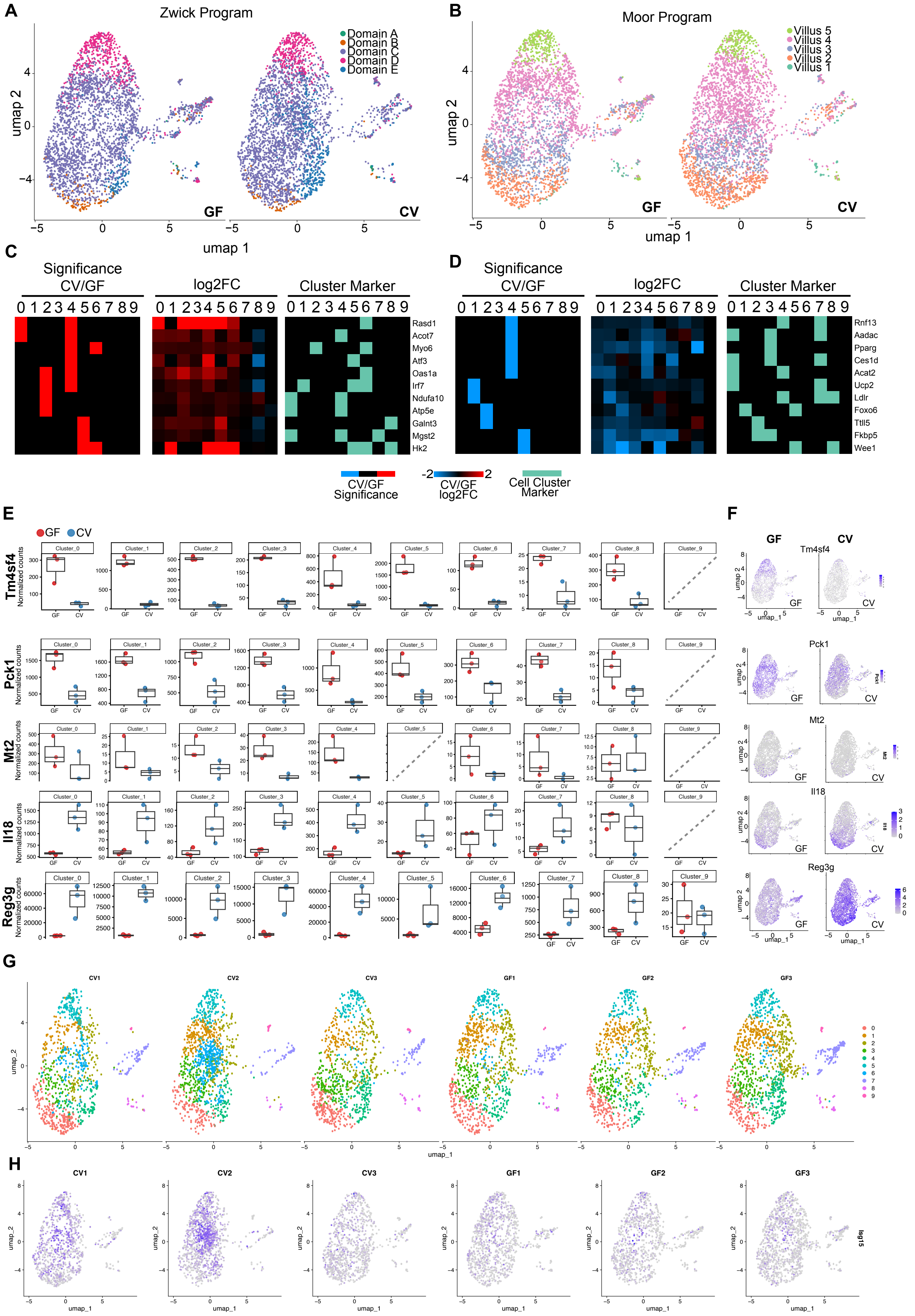
A
Zwick Program
Domain A
Domain B
Domain C
Domain D
Domain E
umap 2
umap 1
GF
CV
B
Moor Program
Villus 5
Villus 4
Villus 3
Villus 2
Villus 1
C
Significance
CV/GF
log2FC
Cluster Marker
0 1 2 3 4 5 6 7 8 9
Rasd1
Acot7
Myo6
Atf3
Oas1a
Irf7
Ndufa10
Atp5e
Galnt3
Mgst2
Hk2
D
Rnf13
Aadac
Pparg
Ces1d
Acat2
Ucp2
Ldlr
Foxo6
Ttll5
Fkbp5
Wee1
CV/GF
Significance
-2
2
CV/GF
log2FC
Cell Cluster
Marker
E
GF
CV
Tm4sf4
Pck1
Mt2
Il18
Reg3g
Normalized counts
Cluster_0
Cluster_1
Cluster_2
Cluster_3
Cluster_4
Cluster_5
Cluster_6
Cluster_7
Cluster_8
Cluster_9
F
Tm4sf4
Pck1
Mt2
Il18
Reg3g
umap_1
umap_2
G
CV1
CV2
CV3
GF1
GF2
GF3
umap_1
umap_2
H
Isg15

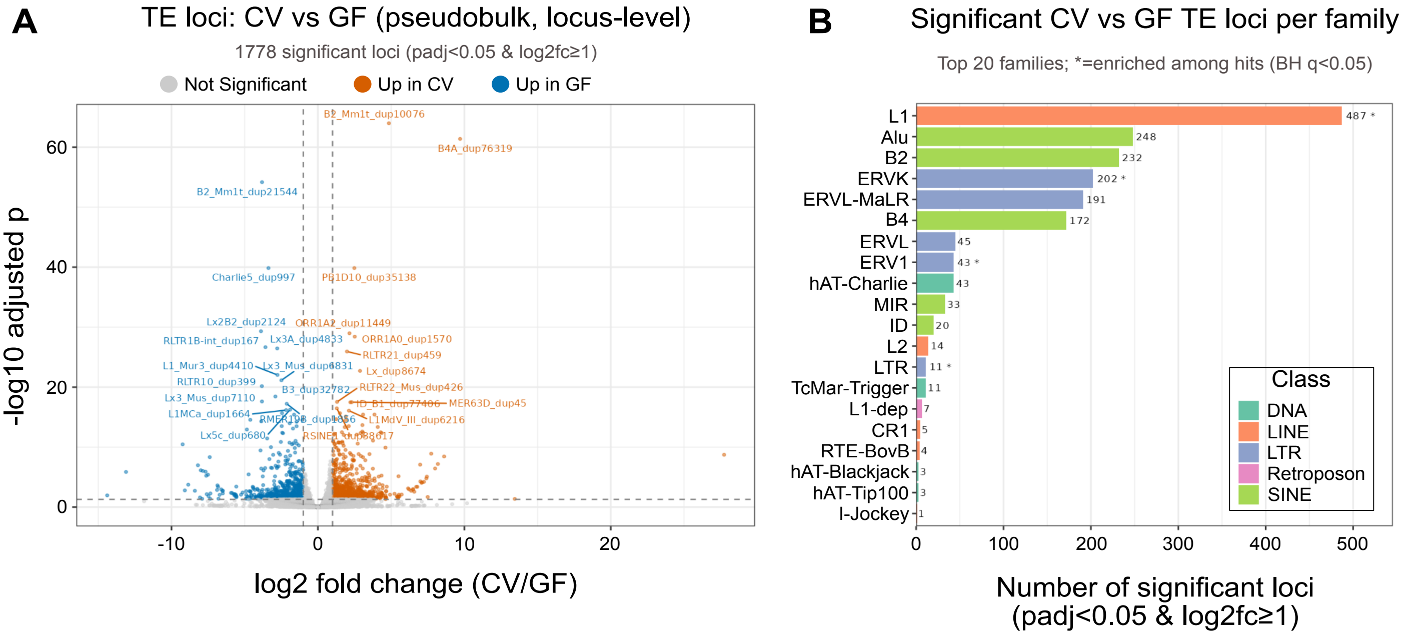
A
TE loci: CV vs GF (pseudobulk, locus-level)
1778 significant loci (padj<0.05 & log2fc≥1)
Not Significant
Up in CV
Up in GF
-log10 adjusted p
log2 fold change (CV/GF)
B2_Mm1t_dup10076
B4A_dup76319
B2_Mm1t_dup21544
Charlie5_dup997
PB1D10_dup35138
Lx2B2_dup2124
ORR1A2_dup11449
RLTR1B-int_dup167
Lx3A_dup4833
ORR1A0_dup1570
RLTR21_dup459
L1_Mur3_dup4410
Lx_dup8674
RLTR10_dup399
B3_dup32782
RLTR22_Mus_dup426
Lx3_Mus_dup7110
MER63D_dup45
L1MCa_dup1664
L1MdV_III_dup6216
Lx5c_dup680
B
Significant CV vs GF TE loci per family
Top 20 families; *=enriched among hits (BH q<0.05)
L1
Alu
B2
ERVK
ERVL-MaLR
B4
ERVL
ERV1
hAT-Charlie
MIR
ID
L2
LTR
TcMar-Trigger
L1-dep
CR1
RTE-BovB
hAT-Blackjack
hAT-Tip100
I-Jockey
487 *
248
232
202 *
191
172
45
43 *
43
33
20
14
11 *
11
7
5
4
3
3
1
Class
DNA
LINE
LTR
Retroposon
SINE
Number of significant loci
(padj<0.05 & log2fc≥1)

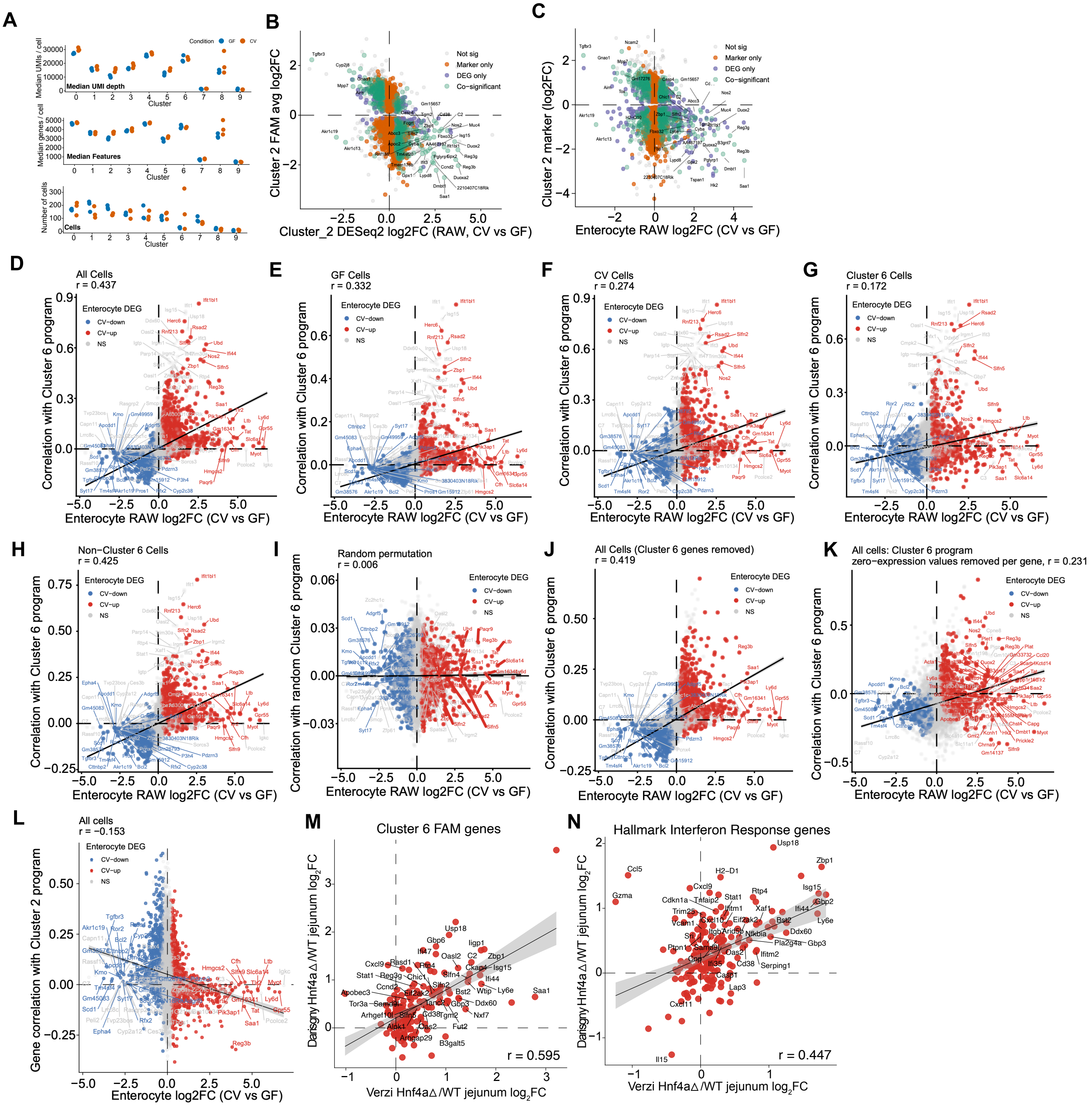
A
Condition GF CV
Median UMI depth
Median UMIs / cell
Median Features
Median genes / cell
Cells
Number of cells
Cluster
B
Not sig
Marker only
DEG only
Co−significant
Cluster 2 FAM avg log2FC
Cluster_2 DESeq2 log2FC (RAW, CV vs GF)
C
Cluster 2 marker (log2FC)
Enterocyte RAW log2FC (CV vs GF)
D
All Cells
r = 0.437
E
GF Cells
r = 0.332
F
CV Cells
r = 0.274
G
Cluster 6 Cells
r = 0.172
Enterocyte DEG
CV−down
CV−up
NS
Correlation with Cluster 6 program
Enterocyte RAW log2FC (CV vs GF)
H
Non−Cluster 6 Cells
r = 0.425
I
Random permutation
r = 0.006
Correlation with random Cluster 6 program
J
All Cells (Cluster 6 genes removed)
r = 0.419
K
All cells: Cluster 6 program
zero−expression values removed per gene, r = 0.231
L
All cells
r = −0.153
Gene correlation with Cluster 2 program
Enterocyte log2FC (CV vs GF)
M
Cluster 6 FAM genes
r = 0.595
Darsigny Hnf4a∆/WT jejunum log2FC
Verzi Hnf4a∆/WT jejunum log2FC
N
Hallmark Interferon Response genes
r = 0.447

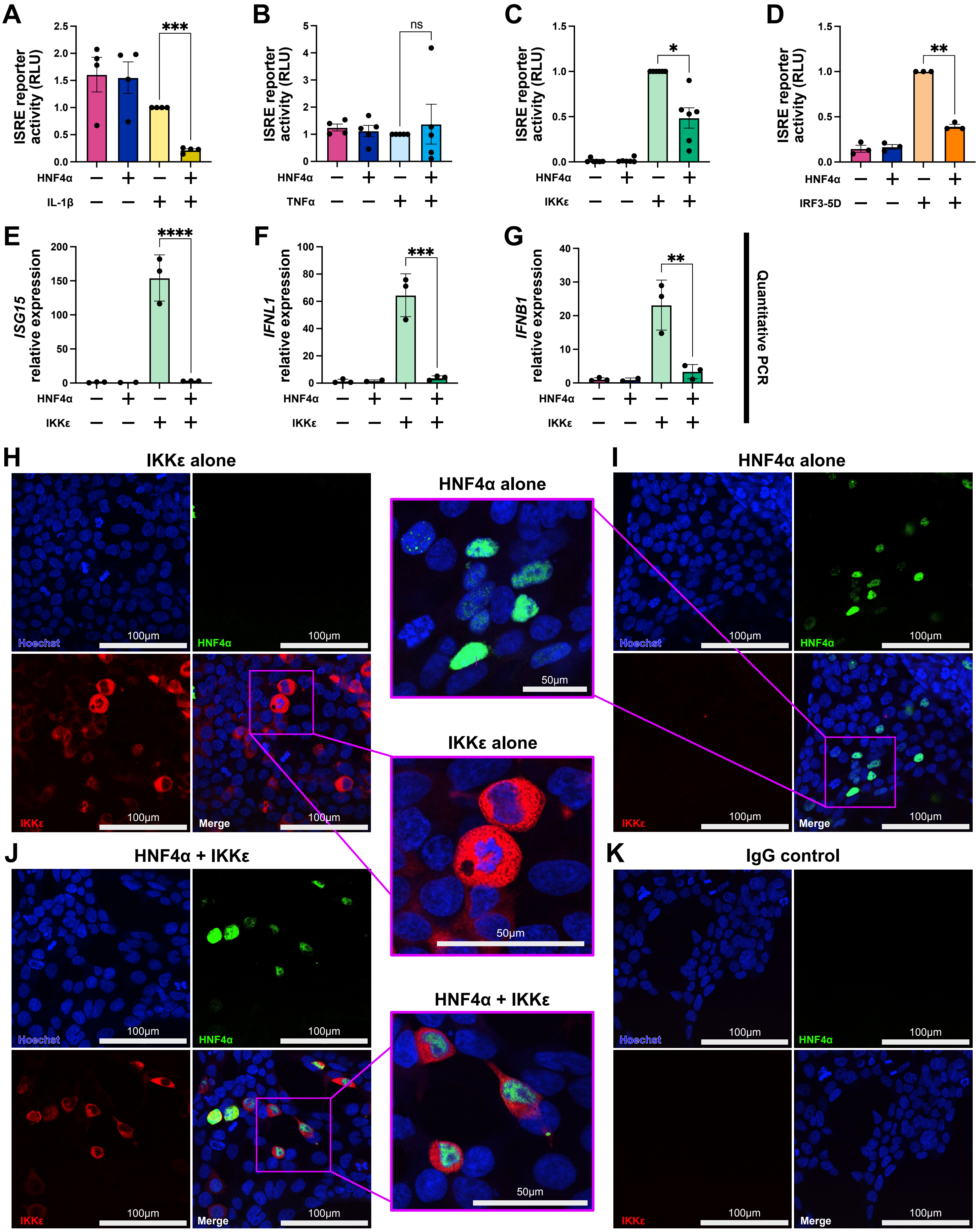

A
ISRE reporter activity (RLU)
***
HNF4α
IL-1β
B
ns
TNFα
C
*
IKKε
D
**
IRF3-5D
E
ISG15 relative expression
****
F
IFNL1 relative expression
***
G
IFNB1 relative expression
**
Quantitative PCR
H
IKKε alone
Hoechst
HNF4α
IKKε
Merge
100µm
50µm
HNF4α alone
IKKε alone
HNF4α + IKKε
I
HNF4α alone
J
HNF4α + IKKε
K
IgG control

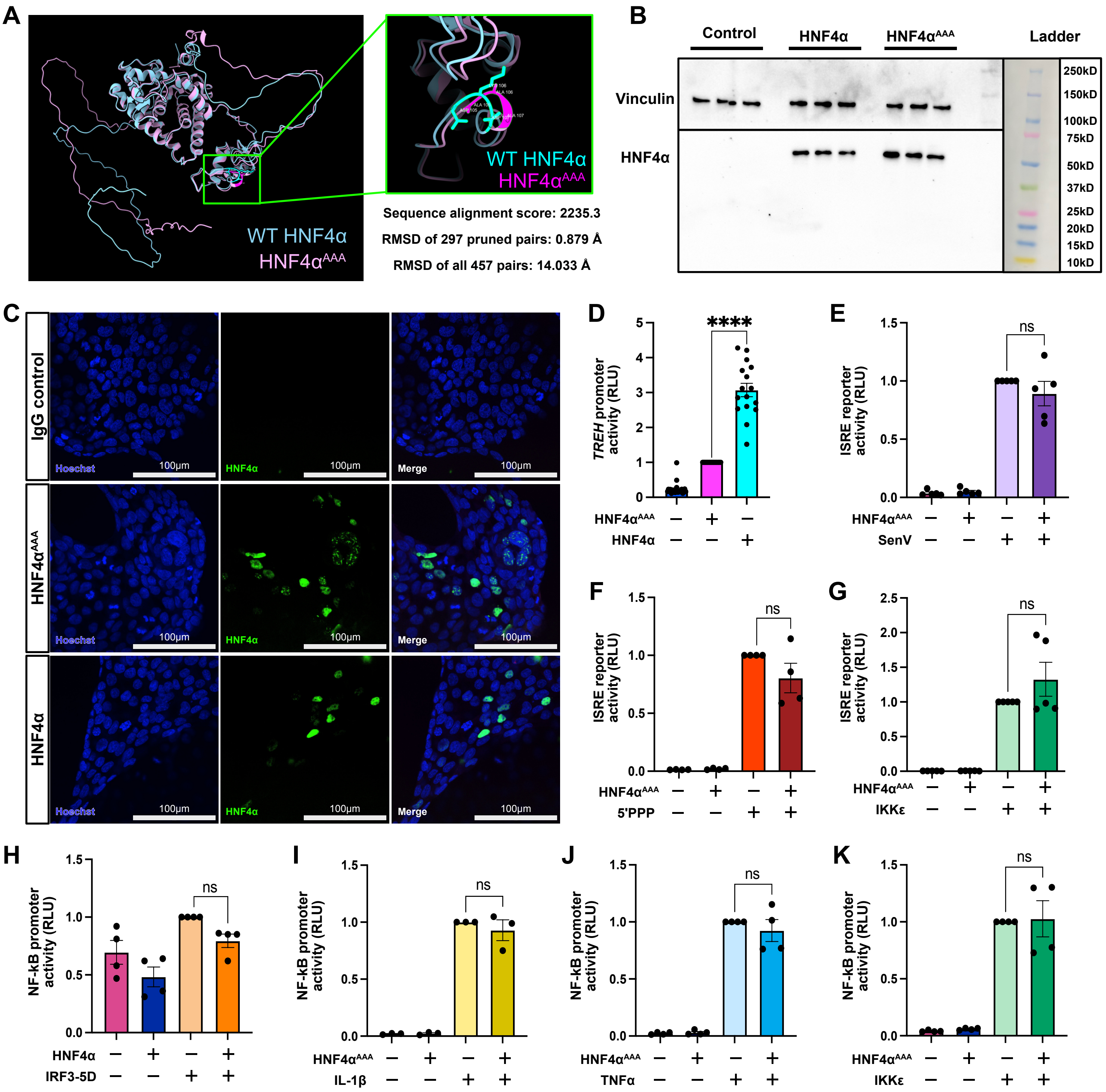
A
WT HNF4α
HNF4αAAA
Sequence alignment score: 2235.3
RMSD of 297 pruned pairs: 0.879 Å
RMSD of all 457 pairs: 14.033 Å
B
Control
HNF4α
HNF4αAAA
Ladder
Vinculin
HNF4α
250kD
150kD
100kD
75kD
50kD
37kD
25kD
20kD
15kD
10kD
C
IgG control
HNF4αAAA
HNF4α
Hoechst
HNF4α
Merge
100μm
D
TREH promoter activity (RLU)
****
HNF4αAAA
HNF4α
E
ISRE reporter activity (RLU)
ns
HNF4αAAA
SenV
F
ISRE reporter activity (RLU)
ns
HNF4αAAA
5'PPP
G
ISRE reporter activity (RLU)
ns
HNF4αAAA
IKKε
H
NF-kB promoter activity (RLU)
ns
HNF4α
IRF3-5D
I
NF-kB promoter activity (RLU)
ns
HNF4αAAA
IL-1β
J
NF-kB promoter activity (RLU)
ns
HNF4αAAA
TNFα
K
NF-kB promoter activity (RLU)
ns
HNF4αAAA
IKKε

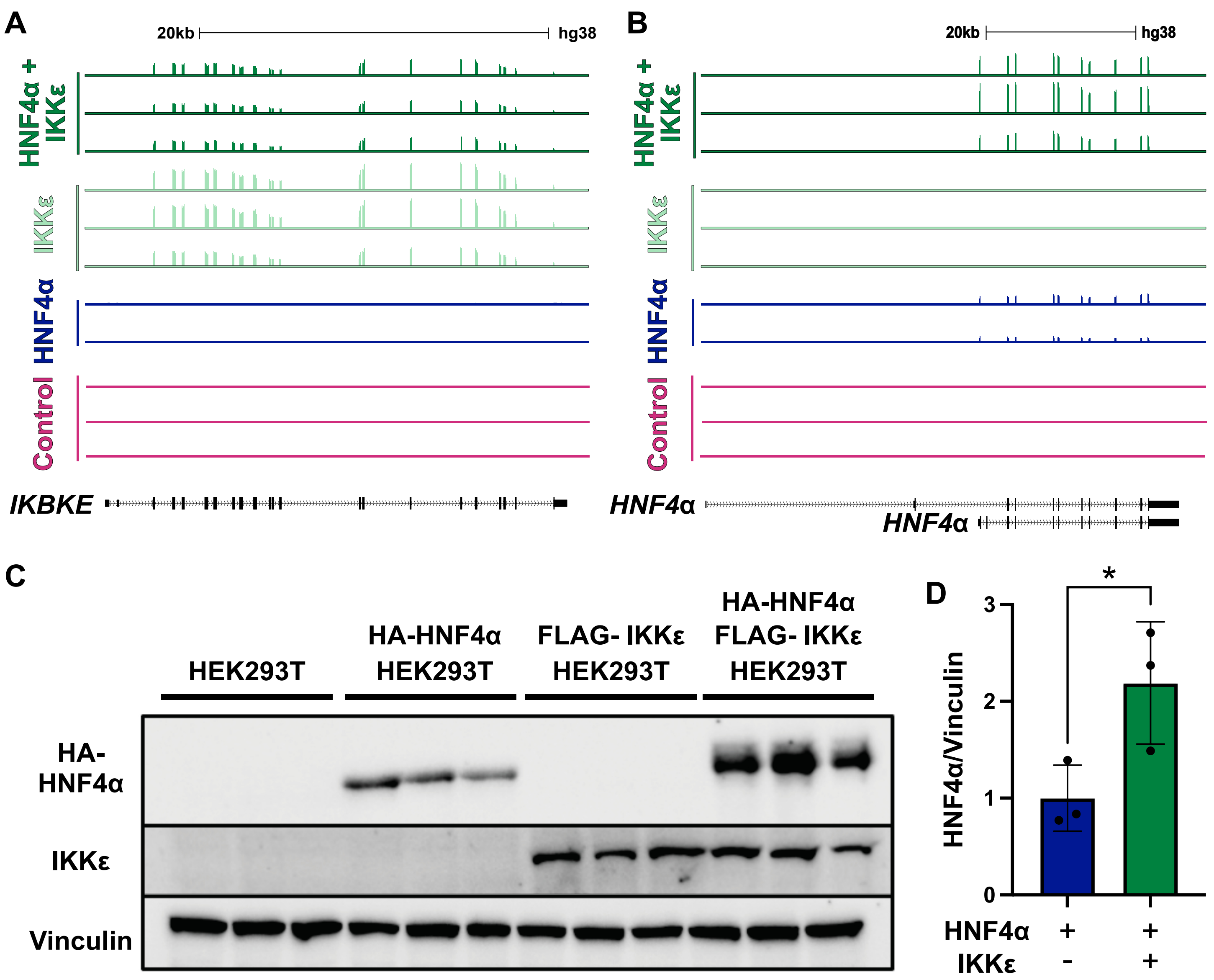

A
B
C
D
20kb
hg38
HNF4α + IKKε
IKKε
HNF4α
Control
IKBKE
HNF4α
HEK293T
HA-HNF4α HEK293T
FLAG- IKKε HEK293T
HA-HNF4α FLAG- IKKε HEK293T
HA-HNF4α
IKKε
Vinculin
HNF4α/Vinculin
*
0
1
2
3
HNF4α + +
IKKε - +

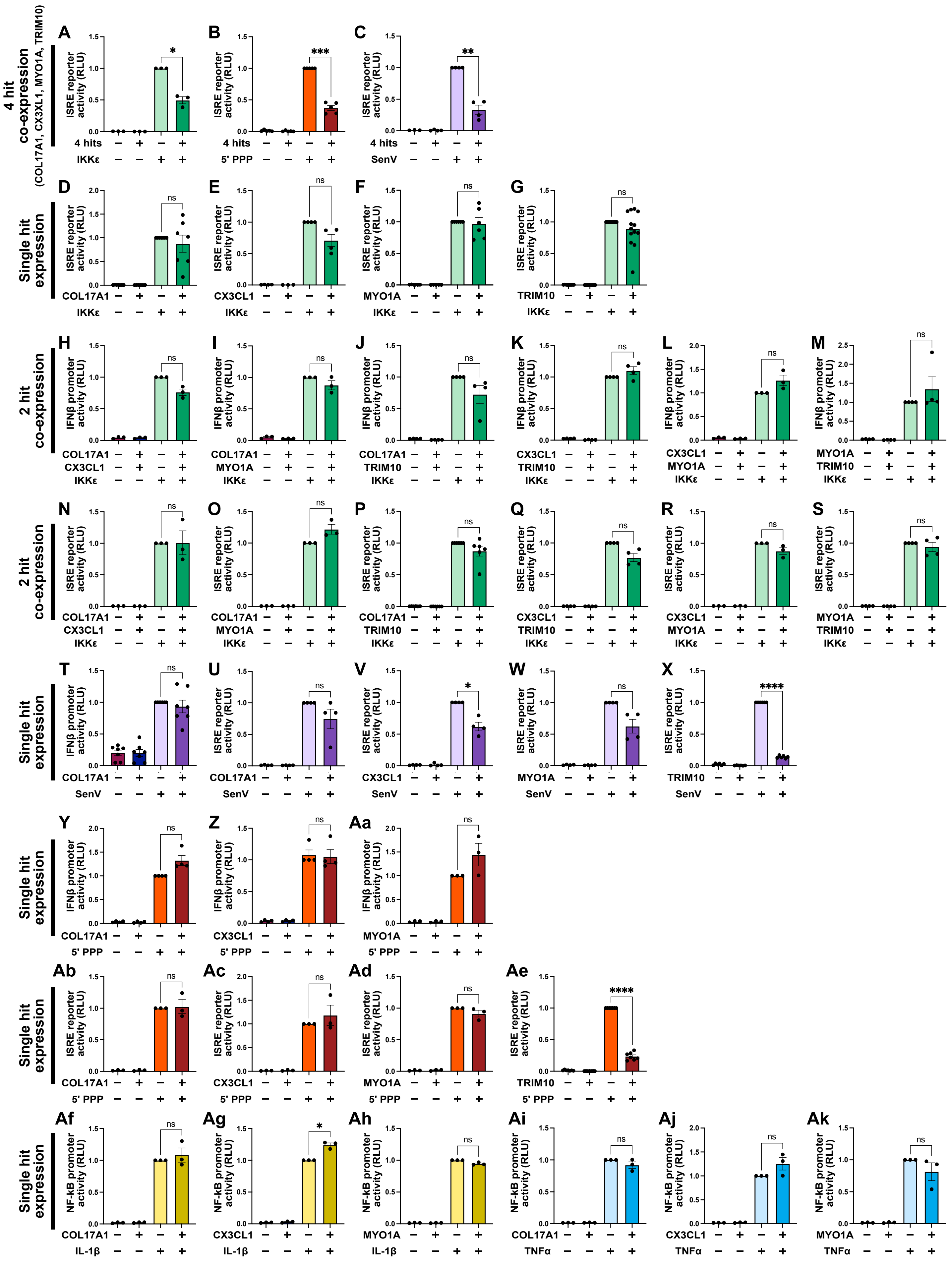

4 hit co-expression
(COL17A1, CX3XL1, MYO1A, TRIM10)
Single hit expression
2 hit co-expression
ISRE reporter activity (RLU)
IFNβ promoter activity (RLU)
NF-kB promoter activity (RLU)
4 hits
IKKε
5' PPP
SenV
COL17A1
CX3CL1
MYO1A
TRIM10
IL-1β
TNFα
ns

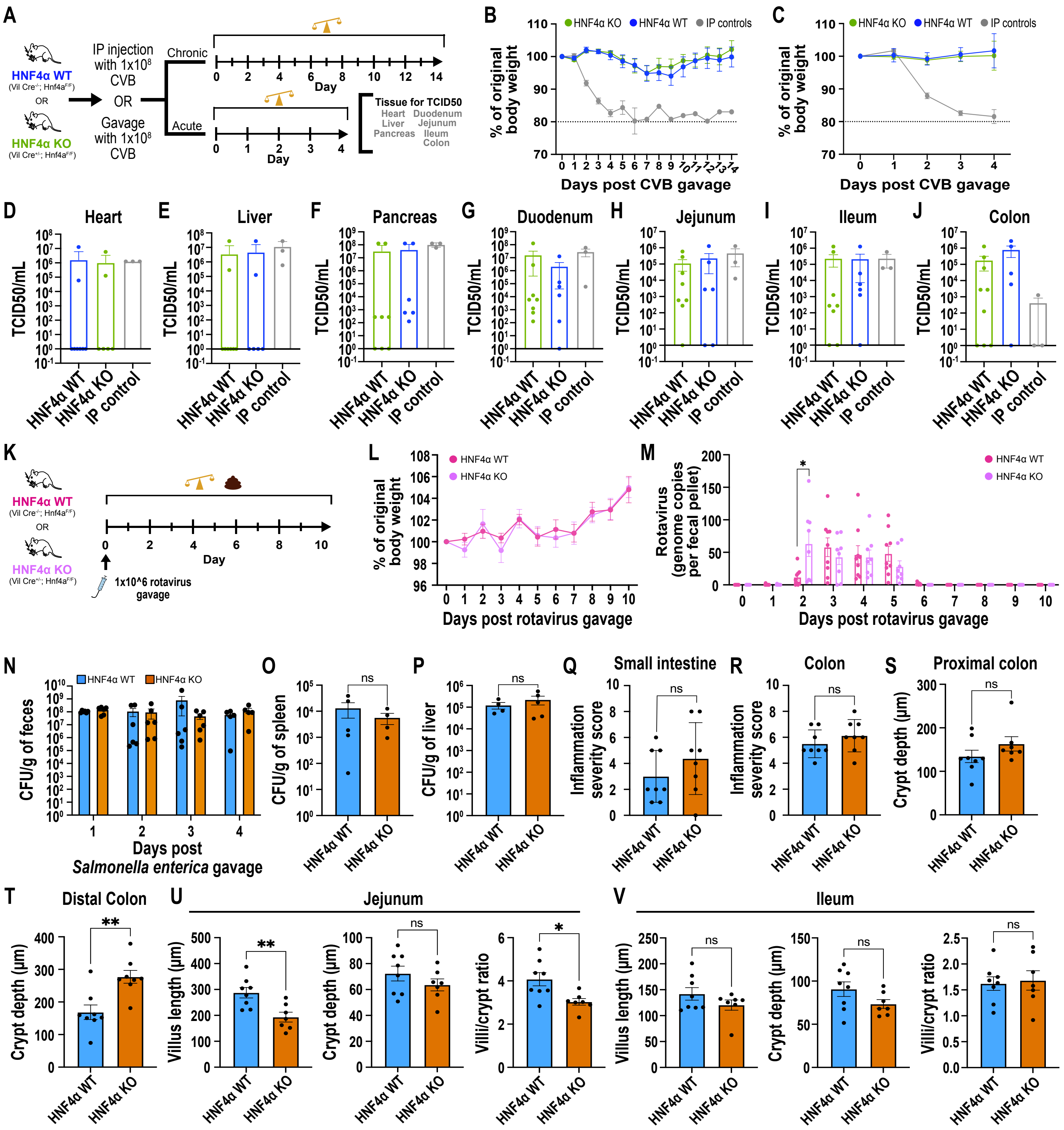

A
HNF4α WT (Vil Cre-/-; Hnf4aF/F)
OR
HNF4α KO (Vil Cre+/-; Hnf4aF/F)
IP injection with 1x10^8 CVB
OR
Gavage with 1x10^8 CVB
Chronic
Acute
Day
Tissue for TCID50
Heart
Liver
Pancreas
Duodenum
Jejunum
Ileum
Colon
B
C
HNF4α KO
HNF4α WT
IP controls
% of original body weight
Days post CVB gavage
D Heart
E Liver
F Pancreas
G Duodenum
H Jejunum
I Ileum
J Colon
TCID50/mL
HNF4α WT
HNF4α KO
IP control
K
1x10^6 rotavirus gavage
L
M
Rotavirus (genome copies per fecal pellet)
Days post rotavirus gavage
N
CFU/g of feces
Days post Salmonella enterica gavage
O
CFU/g of spleen
P
CFU/g of liver
Q Small intestine
R Colon
Inflammation severity score
S Proximal colon
T Distal Colon
Crypt depth (μm)
U Jejunum
V Ileum
Villus length (μm)
Villi/crypt ratio
ns

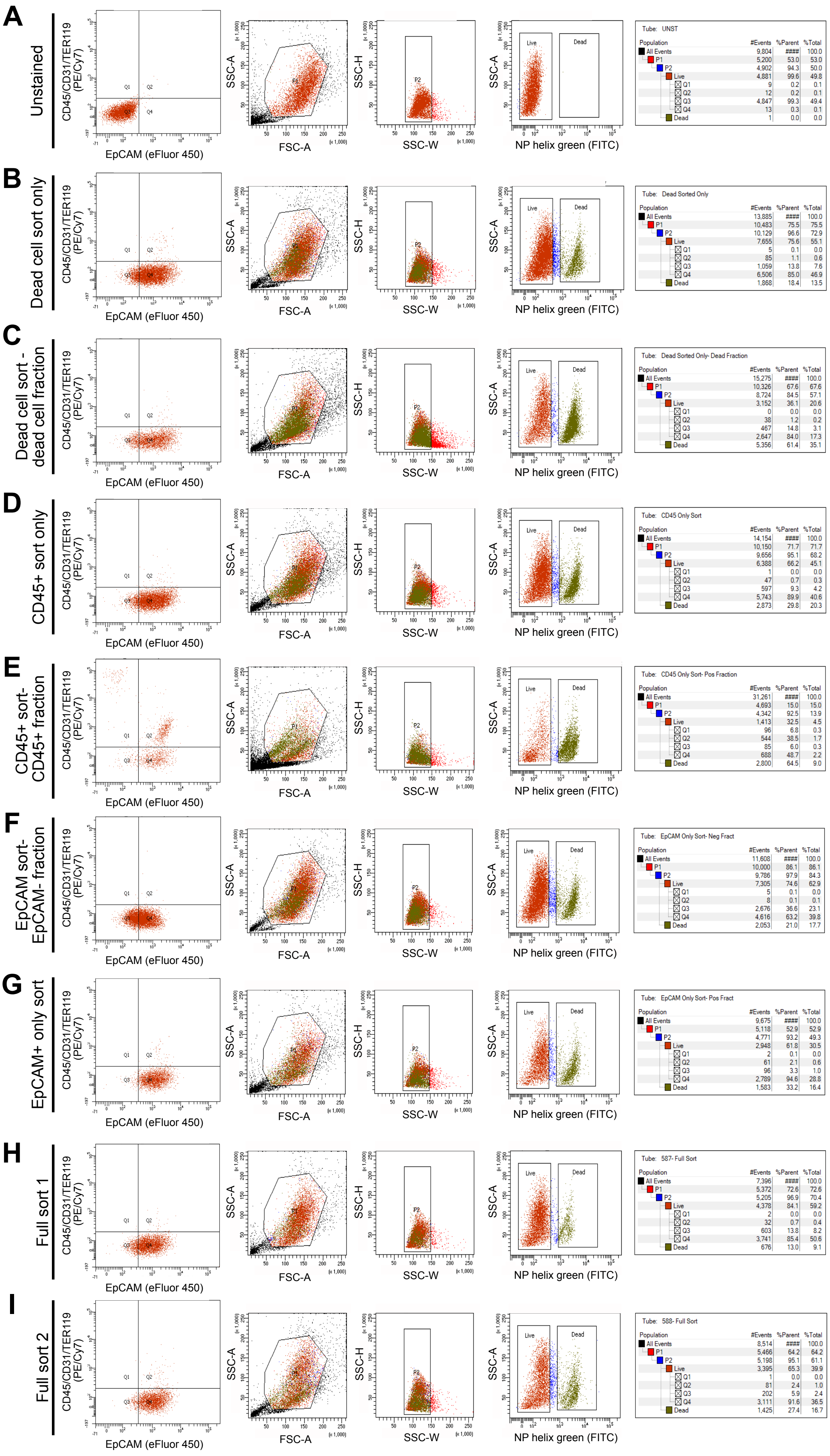

A
Unstained
CD45/CD31/TER119 (PE/Cy7)
EpCAM (eFluor 450)
SSC-A
FSC-A
SSC-H
SSC-W
NP helix green (FITC)
Live
Dead
Tube: UNST
Population #Events %Parent %Total
All Events 9,804 #### 100.0
P1 5,200 53.0 53.0
P2 4,902 94.3 50.0
Live 4,881 99.6 49.8
Q1 9 0.2 0.1
Q2 12 0.2 0.1
Q3 4,847 99.3 49.4
Q4 13 0.3 0.1
Dead 1 0.0 0.0
B
Dead cell sort only
Tube: Dead Sorted Only
Population #Events %Parent %Total
All Events 13,885 #### 100.0
P1 10,483 75.5 75.5
P2 10,129 96.6 72.9
Live 7,655 75.6 55.1
Q1 5 0.1 0.0
Q2 85 1.1 0.6
Q3 1,059 13.8 7.6
Q4 6,506 85.0 46.9
Dead 1,868 18.4 13.5
C
Dead cell sort - dead cell fraction
Tube: Dead Sorted Only- Dead Fraction
Population #Events %Parent %Total
All Events 15,275 #### 100.0
P1 10,326 67.6 67.6
P2 8,724 84.5 57.1
Live 3,152 36.1 20.6
Q1 0 0.0 0.0
Q2 38 1.2 0.2
Q3 467 14.8 3.1
Q4 2,647 84.0 17.3
Dead 5,356 61.4 35.1
D
CD45+ sort only
Tube: CD45 Only Sort
Population #Events %Parent %Total
All Events 14,154 #### 100.0
P1 10,150 71.7 71.7
P2 9,656 95.1 68.2
Live 6,388 66.2 45.1
Q1 1 0.0 0.0
Q2 47 0.7 0.3
Q3 597 9.3 4.2
Q4 5,743 89.9 40.6
Dead 2,873 29.8 20.3
E
CD45+ sort- CD45+ fraction
Tube: CD45 Only Sort- Pos Fraction
Population #Events %Parent %Total
All Events 31,261 #### 100.0
P1 4,693 15.0 15.0
P2 4,342 92.5 13.9
Live 1,413 32.5 4.5
Q1 96 6.8 0.3
Q2 544 38.5 1.7
Q3 85 6.0 0.3
Q4 688 48.7 2.2
Dead 2,800 64.5 9.0
F
EpCAM sort- EpCAM- fraction
Tube: EpCAM Only Sort- Neg Fract
Population #Events %Parent %Total
All Events 11,608 #### 100.0
P1 10,000 86.1 86.1
P2 9,786 97.9 84.3
Live 7,305 74.6 62.9
Q1 5 0.1 0.0
Q2 8 0.1 0.1
Q3 2,676 36.6 23.1
Q4 4,616 63.2 39.8
Dead 2,053 21.0 17.7
G
EpCAM+ only sort
Tube: EpCAM Only Sort- Pos Fract
Population #Events %Parent %Total
All Events 9,675 #### 100.0
P1 5,118 52.9 52.9
P2 4,771 93.2 49.3
Live 2,948 61.8 30.5
Q1 2 0.1 0.0
Q2 61 2.1 0.6
Q3 96 3.3 1.0
Q4 2,789 94.6 28.8
Dead 1,583 33.2 16.4
H
Full sort 1
Tube: 587- Full Sort
Population #Events %Parent %Total
All Events 7,396 #### 100.0
P1 5,372 72.6 72.6
P2 5,205 96.9 70.4
Live 4,378 84.1 59.2
Q1 2 0.0 0.0
Q2 32 0.7 0.4
Q3 603 13.8 8.2
Q4 3,741 85.4 50.6
Dead 676 13.0 9.1
I
Full sort 2
Tube: 588- Full Sort
Population #Events %Parent %Total
All Events 8,514 #### 100.0
P1 5,466 64.2 64.2
P2 5,198 95.1 61.1
Live 3,395 65.3 39.9
Q1 1 0.0 0.0
Q2 81 2.4 1.0
Q3 202 5.9 2.4
Q4 3,111 91.6 36.5
Dead 1,425 27.4 16.7
